# Inter-band connectivity and climate shaped Neanderthal extinction and *Homo sapiens*’ dispersal

**DOI:** 10.1101/2025.09.23.678019

**Authors:** M. Vidal-Cordasco, G Rodríguez-Gómez, A.B Marín-Arroyo

## Abstract

To what extent Neanderthal extinction was triggered by climate change or the arrival and dispersal of *Homo sapiens*, and by which mechanisms, remains unresolved. Here, based on how climate-driven changes affected habitat favourability for primary and secondary consumers and Net Primary Productivity, we estimate carrying capacity, herbivore biomass and intra-guild predation pressure during the time Neanderthals and *H. sapiens* lived in Europe (55-30 ka BP). These spatially explicit estimates were incorporated into an agent-based model simulating human demographic dynamics under various scenarios. Results indicate that carrying capacity changes cannot explain Neanderthal extinction at a continental scale, though they are essential to understanding spatiotemporal distribution patterns of both human species. Conversely, the arrival of *H. sapiens* increased Neanderthal extinction likelihood without requiring a selective advantage. Due to the successive demographic expansions and contractions, group connectivity emerges as the key factor shaping population stability in both species. These findings support that Neanderthal disappearance and *H. sapiens* dispersals were interconnected demographic processes.

---

Extinction is the final step in the evolutionary history of all species^1,2^. In mammals, extinctions are typically multifactorial events, arising from the interplay between intrinsic factors (e.g., life history traits) and extrinsic pressures (e.g., climate change)^2–4^. While anthropogenic pressures dominate extinction drivers nowadays, the causes of Late Pleistocene extinctions remain debated, with ongoing discussions over whether climate change or the arrival and dispersal of *Homo sapiens* was the primary trigger^5–7^.

Among Late Pleistocene extinctions, that of the Neanderthals during Marine Isotope Stage 3 (ca. 60–27 ka BP) remains one of the most contentious issues and is considered a profound upheaval in the history of humanity^8^. Genetic evidence indicates that Neanderthal populations underwent multiple demographic turnovers prior to the arrival of *H. sapiens*^9,10^. However, recent research indicates that small and isolated population groups were not unique to Neanderthals^11,12^. *H. sapiens* arrived and settled in Europe in successive migratory events of small bands that experienced demographic bottlenecks and did not contribute to the ancestry of later European hunter-gatherers^13,14^. In this context, the archaeological record indicates that the replacement of indigenous Neanderthals by *H. sapiens* was neither a single nor a rapid event, but rather a complex and protracted process that gave rise to a biologically and culturally diverse landscape across the continent^15,16^.

In some regions, the Middle to Upper Palaeolithic transition was marked by the emergence of new techno-complexes and limited episodes of genetic admixture between the two human species^17^. Recent studies show that in regions where both human species coexisted, ecosystem productivity was higher or more stable^18^, likely due to local or regional environmental conditions being less sensitive to the global climate oscillations associated with the Dansgaard– Oeschger events^19^ (i.e., alternating cooler stadial and milder interstadial periods)^20–22^. Nonetheless, in most regions, a hiatus between the late Neanderthal and the first *H. sapiens* occupations suggests that Neanderthals disappeared shortly before the arrival of our species, an event often associated with periods of climate deterioration^23–26^. This raises the question of the extent to which the arrival and dispersal of *H. sapiens* and the disappearance of Neanderthals were interconnected demographic processes, and if so, what mechanisms were involved. It also questions the degree to which climate and environmental changes contributed to Neanderthal extinction, and through which specific mechanisms.

To date, most hypotheses on Neanderthal extinction rely on correlative evidence (e.g., timing overlaps between climate change, the arrival of *H. sapiens*, and cultural transitions), but lack explanatory power due to the absence of established causal mechanisms. Consequently, mechanistic rather than correlative models are still necessary for uncovering the causal links underlying these complex dynamics^27^. Agent-based modelling differs from traditional inductive and deductive inference methods by focusing less on prediction and more on exploring the emergent outcomes that arise from imposing a set of conditions, thereby helping to uncover causal mechanisms in complex systems^28,29^. In this study, we propose a three-step modelling framework with the aim of: (1) quantifying the effects of MIS3 climate changes on the spatial distribution of primary and secondary consumer species; (2) assessing the impact of MIS3 climate conditions on the maximum sustainable population densities (i.e., carrying capacity) of *H. neanderthalensis* and *H. sapiens*; and (3) evaluating whether the arrival of *H. sapiens* in Europe increased the likelihood of Neanderthal extinction, and if so, identify the circumstances and mechanisms involved.

First, we used validated paleoclimate simulations alongside a comprehensive database of European archaeological and paleontological MIS3-dated sites to build species distribution models (SDMs) for all primary and secondary consumers weighting over 1 kg. These SDMs quantify how climate fluctuations during MIS3 affected habitat favourability and geographic distribution of species. Next, we integrated the habitat favourability estimates into spatially explicit reconstructions of carrying capacity for both *H. sapiens* and *H. neanderthalensis* at 1,000-year intervals between 55 and 27 kyr BP. This step accounted for bottom-up trophic dynamics influenced by climate-driven fluctuations in Net Primary Productivity and herbivore biomass, as well as intra-guild predation pressure among secondary consumers based on edible biomass, species-specific prey size preferences, and energetic requirements. Lastly, to assess whether and to what extent climate-driven fluctuations in carrying capacity might have influenced Neanderthal extinction, we developed the NEAnderthal Replacement (NEAR) agent-based model. The NEAR model simulates human demographic dynamics across diverse scenarios to assess extinction risk based on four factors: (a) diet composition (percentage of animal-source vs plant-source intake), (b) net population growth rate, (c) demographic stochasticity, and (d) mobility patterns, including logistical, residential, and inter-band mobility. These factors were analysed under two scenarios: (1) in absence of *H. sapiens*, and (2) with the arrival of *H. sapiens* in Europe. In the two-species scenario, additional variables were included to evaluate their effects on Neanderthal extinction risk and timing, such as the number of *H. sapiens* migration waves, population size, probability of interspecific mating, and degree of territorial overlap or segregation between bands.

This study, therefore, tests the following hypotheses:

- H_1_: Climate-driven changes in carrying capacity triggered the extinction of *H. neanderthalensis* in Europe.
- H_2_: The arrival and expansion of *H. sapiens* in Europe had a greater impact on Neanderthal extinction than climate change.

## RESULTS

### Climate-driven transformations

A sustained decline in herbivore biomass began around 45 ky BP, with the lowest values observed at the end of Marine Isotope Stage 3 (MIS3), around 27 kyr BP, and the most pronounced reduction occurring between 45 and 40 kyr BP (Figure 1a). This decline was driven by two main factors: an average 26% decrease in Net Primary Productivity (NPP) (Supplementary Fig. 1), and a contraction in habitat favourability for most herbivore species between 45 and 35 kyr BP. Throughout MIS3, cold-adapted herbivores (e.g., *Coelodonta antiquitatis, Mammuthus primigenius, Ovibos moschatus* or *Rangifer tarandus*) experienced a mean increment of two standard deviations in habitat favourability across Europe, whereas temperate species exhibited a marked decline (Figure 1b). Concurrently, habitat favourability for secondary consumers dropped, on average, by one standard deviation, reaching its lowest values by 35 kyr BP (Figure 1c). As a result, competition among secondary consumers, as indicated by the Global Competition Index, increased threefold from 45 kyr BP until 27 kyr BP (Figure 1 a). These results indicate that climate-driven changes in habitat favourability and herbivore biomass during MIS3 reduced prey availability for most secondary consumers, thereby intensifying intra-guild competition dynamics. Consequently, carrying capacity declined for both *Homo sapiens* and *Homo neanderthalensis* regardless of the proportion of animal-source or plant-source intake (Figure 1d)

**Fig. 1.**
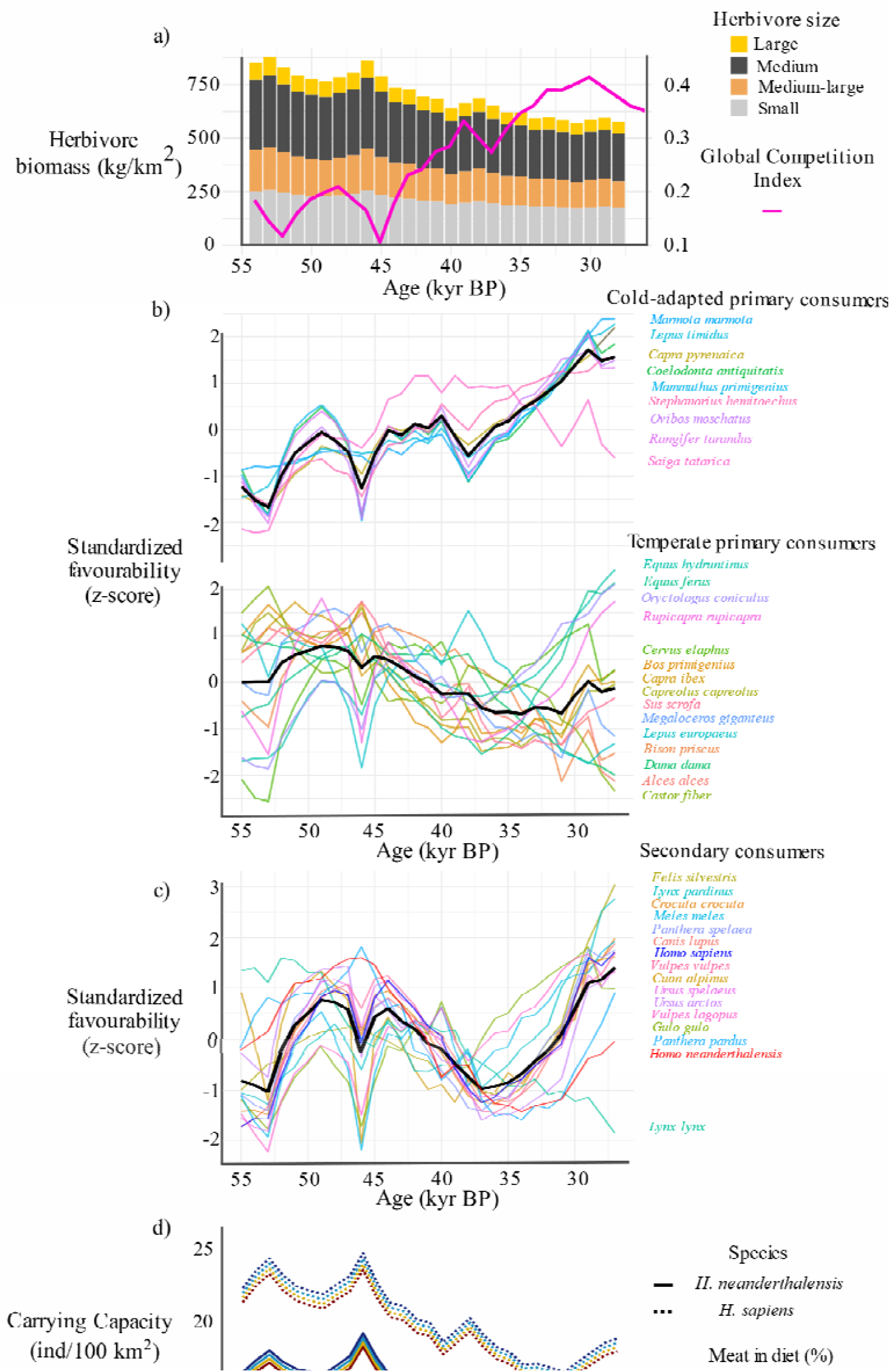
Climate-driven ecological changes. a) Temporal trends in herbivore biomass and the Global Competition Index for secondary consumers; b) Standardized habitat favourability (z-score) over time for cold-adapted (top) and temperate (bottom) herbivores. The bold black line indicates the average value across species within each group; c) Standardized temporal trends in habitat favourability for secondary consumers; d) Human carrying capacity throughout MIS3 under varying assumptions of meat intake.

During MIS3, the mean carrying capacity (CC) for *Homo neanderthalensis* was 14.73 individuals/100 km^2^, whereas *Homo sapiens* exhibited a 25% higher density, averaging 19.83 individuals/100 km^2^ (Figure 2). Although both human species reached their highest CC in southern and western Europe, substantial differences emerged in central and eastern Europe. Specifically, in eastern Balkans, the Anatolian Peninsula, eastern and central Europe, *H. sapiens* displayed a carrying capacity two to four times greater than that of Neanderthals (Figure 2c).

**Fig. 2.**
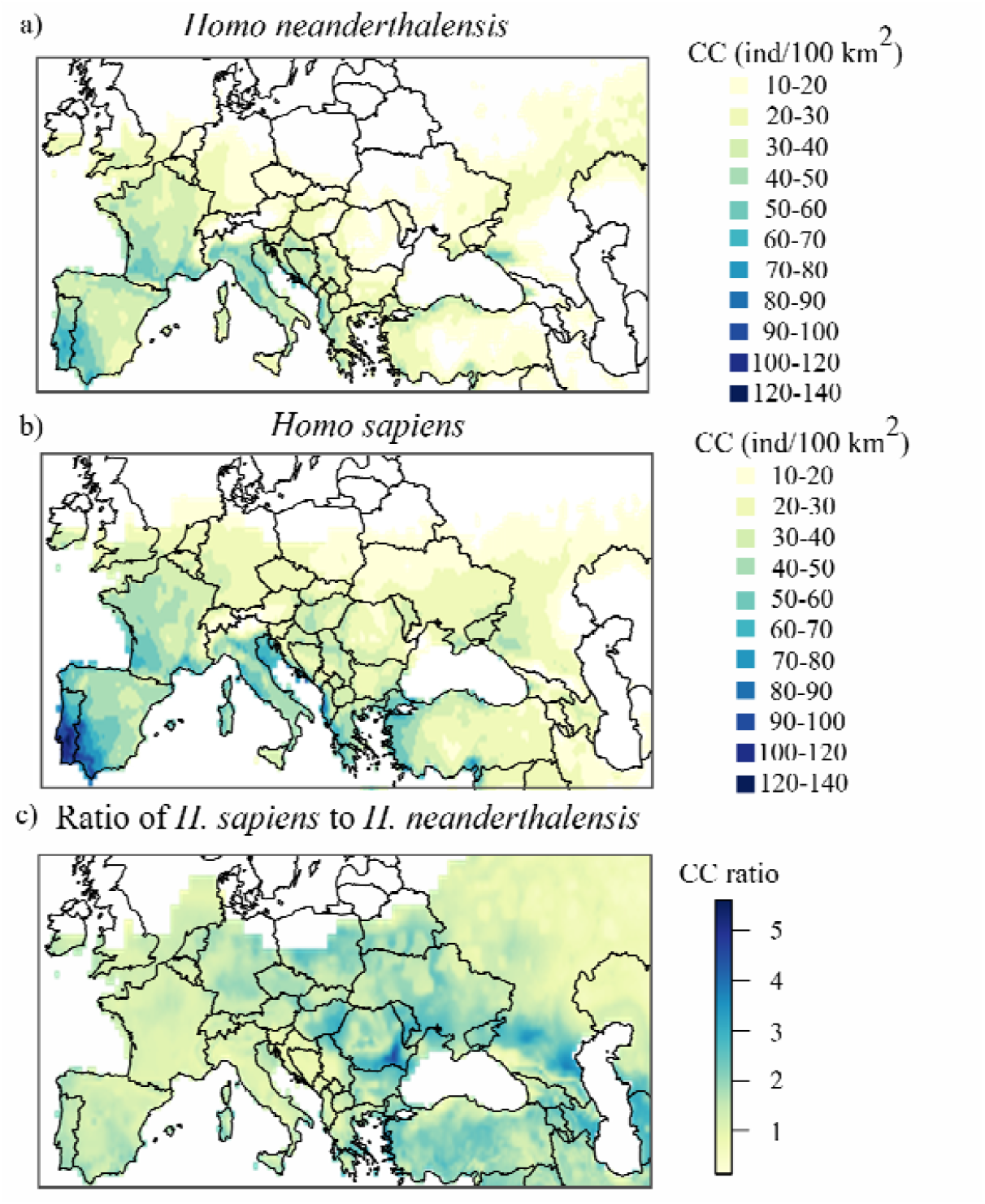
Human carrying capacity during MIS3. a) Mean carrying capacity (Carrying capacity isthe maximum population an environment can sustainably support given the available resources, such as food resources, without causing irreversible environmental degradation.) of Neanderthals in Europe; b) Mean carrying capacity of *Homo sapiens* in Europe; c) Carrying capacity ratio between *H. sapiens* and Neanderthals, expressed as the number of *H. sapiens* per each *H. neanderthalensis*.

### Neanderthal extinction risk

To assess the extent to which the aforementioned climate-driven changes in human carrying capacity may have contributed to Neanderthal extinction, we built an agent-based model (ABM) to evaluate how extinction risk varies in response to four key factors: (a) diet composition (i.e., the proportion of animal-source vs plant-source intake), (b) net population growth rate, (c) demographic stochasticity, and (d) mobility patterns. These variables were analysed under two distinct scenarios: (1) in the absence of *Homo sapiens*, and (2) with the presence of *H. sapiens* in Europe. To quantify the influence of these variables on Neanderthal extinction, we trained a Classification and Regression Tree model on 14,080 simulations (Supplementary Table 1). The CART model achieved an overall accuracy of 89.7%, a precision of 83.6%, and an area under the ROC curve (AUC) of 0.94, indicating strong discriminatory power in distinguishing extinction from survival outcomes based on the ecological and demographic variables.

Results obtained indicate that the arrival and expansion of *H. sapiens* was the most influential factor contributing to Neanderthal extinction (Figure 3). In the single-species scenario, Neanderthals went extinct in 0.5% of simulations (i.e., survived in 99.5% of simulations). However, in the two-species scenario, the extinction rate increased up to 85%. Following the arrival of *H. sapiens*, the primary drivers of Neanderthal extinction risks were: the number of failed *H. sapiens* colonization attempts in Europe, the level of demographic stochasticity affecting both species, and the baseline population growth rate of Neanderthals (Figure 3). These results reject the first hypothesis (H_1_), which posits that climate-driven changes in CC triggered Neanderthal extinction, and instead support the second hypothesis (H_2_), which argues that the arrival and expansion of *H. sapiens* had a greater impact on the likelihood of Neanderthal extinction in Europe during MIS3.

**Fig. 3.**
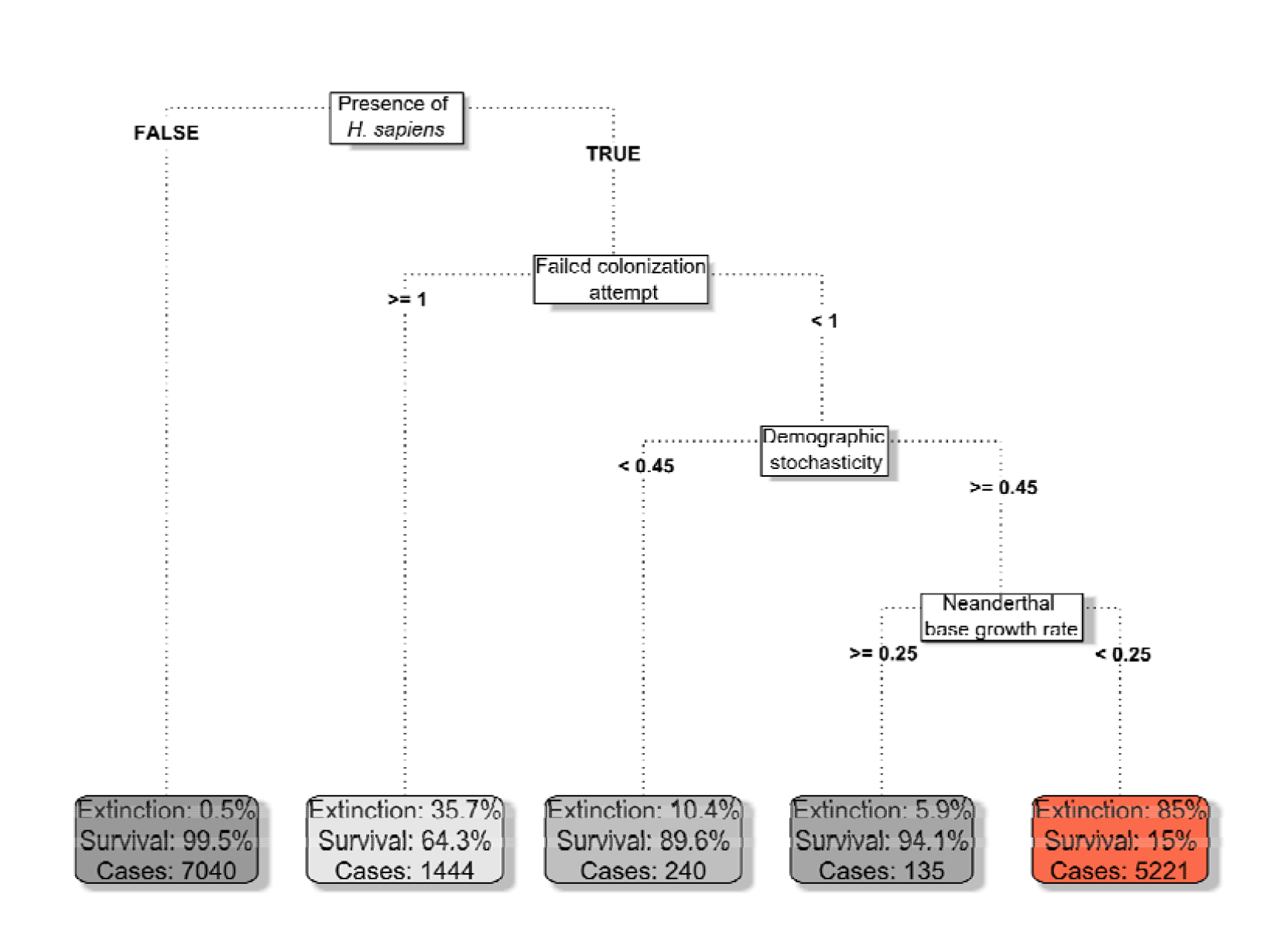
Neanderthal extinction probability. Classification and Regression Tree model showing the main factors contributing to the probability of Neanderthal extinction. The decision tree splits based on key variables, beginning with the presence of *Homo sapiens*, followed by failed colonization attempts, demographic stochasticity, and the Neanderthal baseline growth rate. Each terminal node displays the extinction and survival percentages along with the number of simulated cases falling into that category.

### Timing of Neanderthal extinction

Most of Neanderthal simulated extinction events took place between 51 and 41 kyr BP (Figure 4). However, a number of factors influenced the timing of their extinction. The probability of interbreeding between species, *H. sapiens* baseline growth rates, Neanderthal mating territory size, and the proportion of animal-source intake had no significant effect on the timing of Neanderthal extinction (Figure 4). Conversely, Neanderthal disappearance occurred earlier when their baseline population growth rates were lower than those of *Homo sapiens* and demographic stochasticity was higher (Figure 4). Nonetheless, even when assuming identical demographic patterns for both human species, several factors continued to influence the timing and likelihood of extinction.

**Fig. 4.**
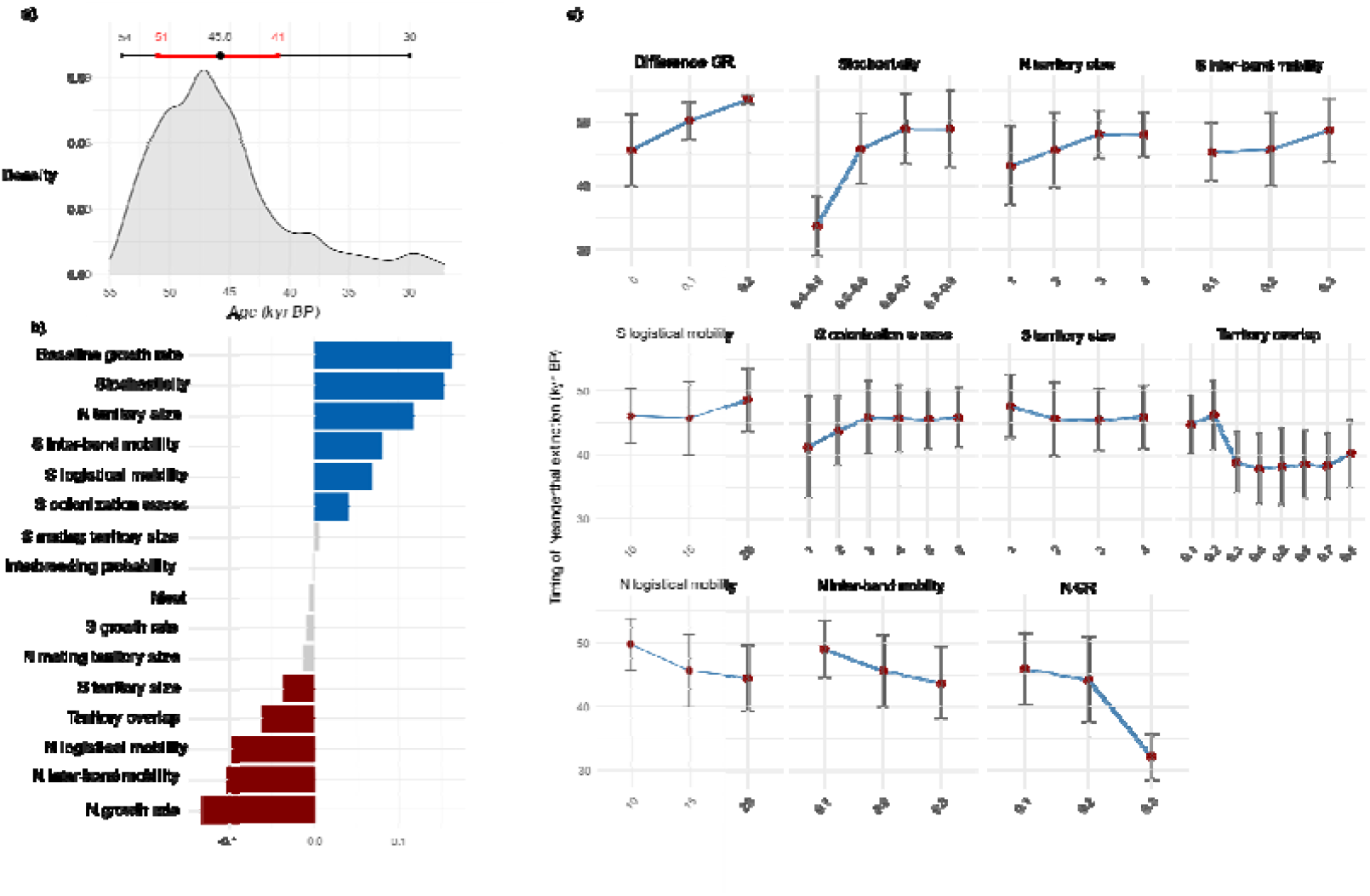
Timing of Neanderthal extinction. a) Density plot showing the simulated timing of Neanderthal disappearance in Europe. Horizontal bars represent the 95.5% (black) and 68.8% (red) confidence intervals across all simulations; b) Spearman correlation coefficients (rho) indicating variables significantly correlated with extinction timing. Blue bars denote significant positive correlations, red bars significant negative correlations, and grey bars uncorrelated variables; c) Mean and standard deviation of extinction timing across different values of each significantly correlated variable (see supplementary Table 1). S= *H. sapiens*, N= *H. neanderthalensis*, GR= baseline growth rate.

Larger Neanderthal territory sizes, as indicated by residential mobility, were associated with earlier extinction, likely due to the increased vulnerability of maintaining sparsely populated regions. Neanderthal territory size showed a significant positive correlation with inter-band distance and a negative correlation with total Neanderthal population size (Figure 5). This suggests that larger territories increased the distance between groups and lowered overall population density, thereby raising the risk of demographic bottlenecks and contributing to earlier demise (Figure 5). In contrast, greater logistical and inter-band mobility were associated with a delayed extinction. Increased logistical mobility facilitated resource exploitation over larger areas, while higher inter-band mobility enhanced population connectivity, promoting gene flow, reducing isolation, and contributing to a longer persistence of Neanderthals in Europe (Figure 4).

**Fig. 5.**
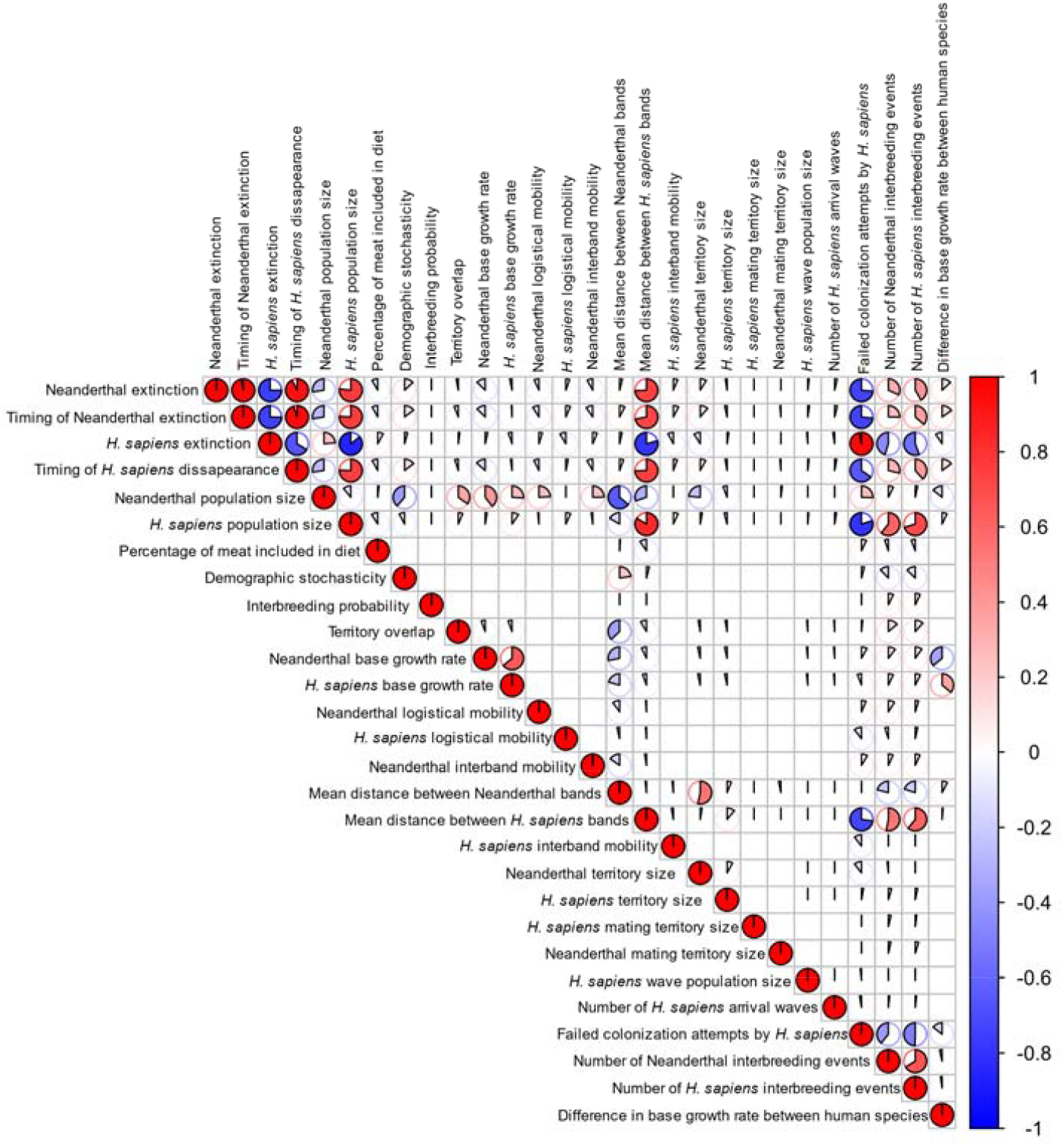
Correlation matrix of variables included in the NEAR model. Pairwise Spearman correlation coefficients among all variables used in the analysis. The matrix visualizes the strength and direction of monotonic relationships, with blue indicating negative correlations and red indicating positive correlations. The size and colour intensity of each pie segment correspond to the magnitude of the correlation coefficient, as shown by the colour scale bar.

Greater logistical and inter-band mobility among *H. sapiens* groups was associated with an earlier Neanderthal extinction, likely due to their effects on accelerating *H. sapiens* expansion into Neanderthal-occupied areas. However, the number of *H. sapiens* colonization waves also influenced extinction timing: as the number of colonization attempts increased, up to a threshold of three waves, the extinction of Neanderthals was delayed (Figure 4). This suggests that early colonization attempts by *H. sapiens* that failed to establish lasting populations might be crucial for the longer persistence of *H. neanderthalensis* in some regions.

Decreased territorial overlap between bands was associated with an earlier Neanderthal extinction, particularly when overlap fell below a threshold of 30% (Fig. 4). In this context, greater spatial overlap, indicating lower segregation, was negatively correlated with the distance between Neanderthal bands and positively correlated with both the frequency of interbreeding events and overall Neanderthal population size (Fig. 5). These findings suggest that, although territorial overlap between bands may have reduced resource availability, it promoted demographic stability by decreasing inter-band distances and facilitating reproduction between groups.

### Replacement patterns

Despite *H. sapiens* spread rapidly toward western longitudes, their population densities remain higher in eastern Europe until approximately 45–43 kyr BP, when their presence markedly increases in central and western Europe (Fig. 6). As *H. sapiens* expanded from east to west, Neanderthal populations progressively retreated westward. However, in the westernmost part of the Balkan Peninsula, Neanderthals persisted until around 40 kya, likely due to higher habitat favourability and carrying capacity in that region. The spatial overlap between the two species gradually increased, becoming complete by 43 ky BP.

**Fig. 6.**
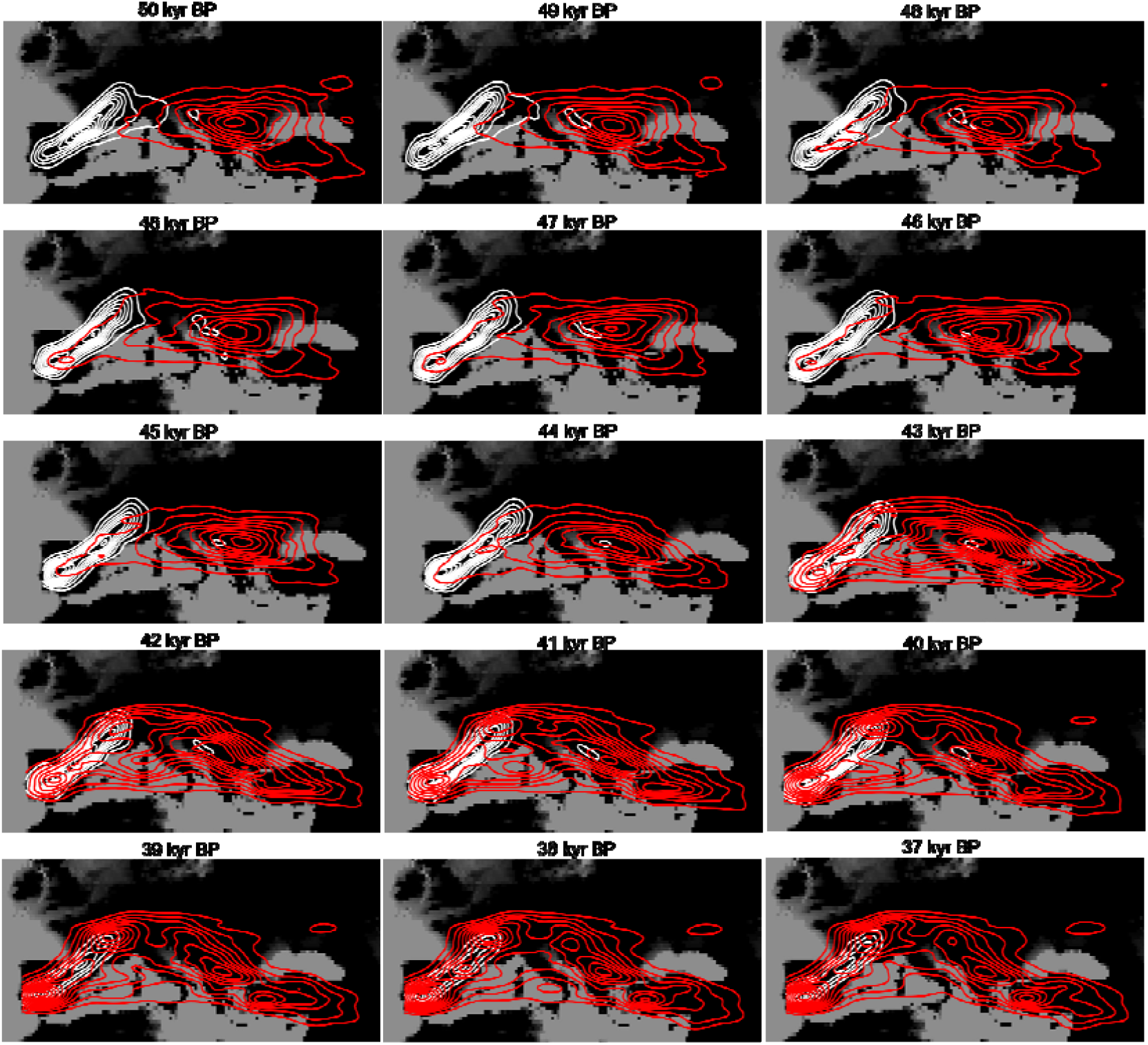
Spatial distribution of agents. Geographic distribution of Neanderthal (white contours) and *H. sapiens* (red contours) populations across Europe during the Middle to Upper Palaeolithic transition according to the NEAR model. Density contours represent kernel density estimates of the mean modelled geographic locations for each species per each 1 kyr years between 50 and 37 kyr BP.

In 66.7% of the simulations, *Homo sapiens* colonization failed at least once, meaning that in most scenarios, *H. sapiens* disappeared from Europe prior to Neanderthals. The number of failed colonization attempts by *H. sapiens* was positively correlated with Neanderthal population size and negatively correlated with both *H. sapiens* population size and their inter-band distance (Figure 5). These findings suggest that *H. sapiens* group sizes, which influenced inter-band distances, along with Neanderthal population densities, influenced the likelihood of *H. sapiens* populations establishing and persisting over time in Europe.

## DISCUSSION

Despite substantial scientific investment and major advances in reconstructing past climate changes^21,30,31^, the specific mechanisms through which these shifts affected terrestrial ecosystems and human populations remain poorly understood. In this study, we presented a palaeoecological modelling framework to reconstruct the effects of climate change on the spatiotemporal dynamics of resource availability and abundance, habitat favourability and intra-guild predatory pressure, key ecological constraints on population sizes. By integrating estimates of human carrying capacity into the NEAR agent-based model, we simulated demographic dynamics under a wide range of scenarios. These results have allowed us to disentangle the respective roles of climate and the arrival of *Homo sapiens* in the extinction of Neanderthals.

MIS3 paleoclimate reconstructions indicate a general decline in mean annual temperatures and precipitations in Europe^32–34^. This study shows that, as a consequence, cold-adapted herbivores experienced a marked increase in habitat favourability, and therefore an expansion of their potential geographic ranges. This is consistent with the archaeological and paleontological record, which indicates a major southward of cold- and dry-adapted species associated with steppe and tundra environments during MIS3^35–38^. In contrast, most temperate herbivores experienced a decline in habitat favourability. Since the cold-adapted faunas, commonly referred to as the *Mammuthus*–*Coelodonta* Faunal Complex^38,39^, included the largest terrestrial mammals of MIS3, such shifts in habitat favourability directly affected prey availability for secondary consumers. As large-bodied herbivores generally fall outside the preferred prey size range of most secondary consumers^40,41^, top-down predation pressure on small and medium-sized herbivores intensified. In addition, declining temperatures and precipitation led to a contraction at the base of the food chain (i.e., NPP), further reducing the available herbivore biomass and intensifying competition among secondary consumers.

Recent studies indicate that, during the Middle to Upper Palaeolithic transition, carnivore co-occurrence and taxonomic diversity increased in both archaeological and paleontological layers across Europe^42^. This pattern is attributed to contractions in habitat favourability, which concentrated secondary consumers, including humans, into smaller and more fragmented areas^42–45^. Results obtained in this study provide additional evidence for an increment in competition among secondary consumers, as reflected by the Global Competition Index. This intensification of competition arose not only from shrinking geographic ranges, but also from a substantial reduction in total available biomass. This biomass reduction is supported by spores of coprophilous fungi, which are commonly used as a proxy for herbivore biomass in sediment cores^46^, and indicate a decline between 44 and 39.8 kya in central Europe^47,48^. As a result, between 45 and 40 kyr BP, the mean carrying capacity for both *Homo sapiens* and *Homo neanderthalensis* declined.

The mean carrying capacity (CC) for both *Homo sapiens* and *Homo neanderthalensis* in Europe falls within the interquartile range of observed densities in extant hunter-gatherer societies^49,50 50^(Supplementary Fig. 2). These values exceed previous density estimates for both Neanderthals^51^ and *H. sapiens*^52,53^ during the Late Pleistocene, despite showing similar spatial patterns^52,53^. Nevertheless, it is important to note that these CC estimates represent environmental population ceilings rather than actual population sizes. In other words, they represent the maximum number of individuals the environment could support, based on habitat favourability, energetic requirements, dietary preferences, the availability and abundance of resources, and under the assumption that all areas are occupied up to their maximum sustainable density. In contrast, when incorporated into the NEAR model to simulate human demographic dynamics, the resulting population size was, on average, 1,916 (S.D. 1,319) individuals for Neanderthals and 3,133 (S.D. 1,768) for *H. sapiens*. These values are notably lower due to three factors incorporated into the NEAR model: (1) not all habitable areas are continuously occupied; (2) extensive regions were uninhabitable during MIS3 due to glacial coverage, represented in the model by areas of low habitat favourability; and (3) human population sizes do not necessarily reach their CC. Both CC and simulated population densities of Neanderthals and *H. sapiens* decreased throughout MIS3; yet, it remains to be tested whether this climate-driven decline was sufficient to drive Neanderthal extinction.

The observed declines in CC contributed to the reduction of the Neanderthal metapopulation and their disappearance from some regions. Specifically, simulations excluding *H. sapiens* showed *H. neanderthalensis* undergoing demographic decline along general east-to-west and north-to-south clines, opposite to the directions followed by *H. sapiens* during their dispersal across Europe, as indicated by the archaeological record^54,55^. This supports the idea that regional Neanderthal depopulation may have facilitated a staggered repopulation by modern humans^23^. However, complete Neanderthal extinction occurred in less than 1% of these simulations. These results suggest that, while climate-driven ecological changes can explain the regional disappearance of Neanderthals and their contraction toward western and southern regions, they do not fully account for their extinction at the continental scale.

When simulations included the presence of *H. sapiens*, the Neanderthal extinction risk increased significantly, even without assuming any difference in baseline population growth rates or mobility patterns between the two species. Additionally, a higher number of failed *H. sapiens* colonization attempts was associated with lower extinction risks for Neanderthals. Drawing inspiration from the concept of neutral allelic fixation through genetic drift, the “species drift hypothesis” proposes that neutral stochastic fluctuations in human demographic dynamics can, over time, account for the replacement of Neanderthals by *H. sapiens*^56^. The results of this study further support this idea, demonstrating that no selective advantage of *H. sapiens* over *H. neanderthalensis* is required to explain their replacement. Nevertheless, the impact of *Homo sapiens’* presence on Neanderthals’ extinction risks and timing depended on multiple specific factors.

The presence of *H. sapiens* in Europe intensified indirect competition for resources, particularly in regions with low or fluctuating trophic resources^18^. However, maintaining sparsely populated bands may have been an even more critical factor contributing to the vulnerability of human populations. The simulations conducted in this study show that greater spatial overlap between bands was negatively correlated with Neanderthal isolation extent and positively correlated with both the frequency of interspecific admixture events and overall Neanderthal population size. Moreover, higher inter-band mobility within or between species was associated with longer populations persistence. This explains why larger territorial overlaps, despite intensifying actual pressure on resources, also facilitated movement and connectivity between bands, promoting gene flow, reducing isolation, and ultimately supporting longer population survival.

Palaeolithic human demography has been characterised by a set of “booms and busts”^57–59^, where climate change, environmental shifts, and human adaptations influenced multiple population turnovers^14,60,61^. In the context of MIS3, this study provides additional evidence suggesting that demographic bottlenecks were the rule rather than the exception^28^, and that connectivity between bands was one of the most influential factors on population stability. Within this dynamic landscape of successive demographic expansions and contractions, the arrival of *H. sapiens* in Europe may have disrupted the potential for Neanderthal populations to re-expand northward and eastward following climate-induced changes in CC. In contrast, the higher *H. sapiens* CC throughout natural dispersal routes in eastern and central Europe could have facilitated successive arrival waves that, in turn, were crucial for inter-group connectivity and repopulation.

One of the most striking features of the Middle to Upper Palaeolithic transition is the regional variability in the timing and patterns of early *H. sapiens* dispersals and the extinction of *H. neanderthalensis*^62–64^. In some areas, new “transitional” cultures emerged^55,65–67^; in others, Middle Palaeolithic techno-complexes disappeared before the arrival of those associated with *Homo sapiens*^18,68^; and elsewhere, genetic evidence suggests admixture between the two species^69,70^. Within this complex context, regional segregation between Neanderthals and *H. sapiens* has been proposed. The Ebro frontier hypothesis suggested that, in the Iberian

Peninsula, Neanderthals were confined to southern latitudes, while *H. sapiens* occupied territories north of the Ebro basin^71,72^ (but see ^73,74^). Likewise, it has been proposed that in the Balkans, Neanderthals persisted in central and western mountainous marginal areas, while *H. sapiens* settled in eastern areas and along river corridors^75^. Our study shows that, although the Ebro basin did not represent an ecological barrier causing differing carrying capacities for the two species, in eastern Balkans, the Danube basin, and central Europe, the CC for *H. sapiens* was two to four times higher than that for Neanderthals (Fig. 2). These results suggest that the higher *H. sapiens* CC along natural dispersal routes (e.g., lower and upper Danube) favoured their rapid spread across the continent. Yet, such dispersals, facilitated by high CC and low Neanderthal densities, did not ensure the long-term establishment of *H. sapiens* populations.

Recent years have witnessed growing interest in the mechanisms underlying *H. sapiens* dispersals and the factors driving the variable success of their expansions into new regions^76–80^. Although the role of climate as a gatekeeper for the initial *H. sapiens* dispersals out of Africa during MIS5 (126–74 ka BP) is well established^28,53,81,82^, it remains unclear whether human or environmental factors influenced their later and intermittent presence in Europe^83,84^, particularly when compared to the rapid dispersals observed in regions such as southern Asia^81,85^. Worsening climate is often cited as a driver of the retreat of early populations back into Africa^86^. In addition, it has been proposed that early *H. sapiens* migrations occurred at low population densities (<5 individuals/100 km^2^), making it likely that these groups were assimilated by prevalent Neanderthal populations, whereas higher population densities from around 45 kyr BP may have favoured more sustained occupations^53^. On the other hand, some authors claim that MIS3 *H. sapiens* populations exhibited not only “anatomical modernity” but also “derived behaviours”^87^ (but see ^88^), potentially enabling them to outcompete Neanderthals^89–91^. The results of this study suggest that the establishment of stable and lasting populations of *H. sapiens* that could replace Neanderthals during MIS3 was contingent on population sizes and the arrival of successive migratory waves.

This study finds no correlation between Neanderthal and *H. sapiens* population sizes, suggesting that Neanderthals did not act as a demographic barrier to *H. sapiens* expansion. However, this result may be influenced by the fact that, by the time *H. sapiens* arrived in Europe, Neanderthals were already marginally present in eastern regions, with their core populations having contracted toward southern and western areas. On the other hand, larger *H. sapiens* populations were associated with higher levels of interspecific admixture events and shorter inter-band distances. These results indicate that it was *H. sapiens* inter-band connectivity, rather than Neanderthal presence or environmental carrying capacity, that played a central role in the establishment of stable *H. sapiens* populations in Europe. The close kinship observed among individuals in archaeological sites suggests that these early groups were small^13,92^. In this connection, genetic evidence indicates that isolation was common among both *H. neanderthalensis*^12^ and *H. sapiens*^13^, with the latter not contributing to the ancestry of later European hunter-gatherers^13^. This lack of genetic continuity, along with small population sizes, is consistent with our results, which indicate that in most cases, early waves of *H. sapiens* disappeared from Europe prior to the complete extinction of Neanderthals.

As proposed by previous studies^28,93,94^, the extinction of Neanderthals was likely a multifactorial process resulting from a complex interplay of extrinsic and intrinsic factors. The results of this study provide additional insight into the underlying mechanisms of this phenomenon. First, the findings show that climate-driven changes in human carrying capacity cannot account for the continental-scale extinction of Neanderthals, although such changes remain essential for understanding the spatiotemporal distribution patterns. In contrast, the arrival of *H. sapiens* played a decisive role in the final extinction of Neanderthals, likely by disrupting their potential to re-expand northward and eastward in response to climate-induced changes in carrying capacity. Nevertheless, no selective advantage of *H. sapiens* over Neanderthals is required to explain their extinction. Rather, connectivity between groups emerges as a critical factor influencing population stability and persistence. In this context, intermittent coexistence and episodes of gene flow between both human species may have enhanced inter-band connectivity, thereby supporting larger and more stable populations. These findings lend further support to the view that the disappearance of Neanderthals and the spread of *H. sapiens* in Europe were not isolated events, but deeply interconnected demographic processes.

## METHODS

### -Database

This study draws on a comprehensive database focused on the European Marine Isotope Stage (MIS) 3^18,20^ and available atosf.io/4aqjk. The dataset includes faunal records from European archaeological and paleontological sites, documenting the presence of herbivore and carnivore species in dated layers. For each assemblage, the database captures taxonomic identifications, cultural affiliations, and associated chronometric determinations. While genus-level identifications were noted, only species-level records were retained for analytical purposes in this study.

Bayesian age modelling was carried out using ChronoModel, where each “event” corresponded to a single chronometric measurement, and multiple determinations from a single sample were grouped as one event. Each “phase” represented a set of events from a distinct stratigraphic layer. Following previous studies, we used the default Markov chain Monte Carlo settings, consisting of three chains, 1000 burn iterations, and 500 batch iterations with a maximum of 20 batches and 100,000 acquisition iterations with a thinning interval of 10^95^. Radiocarbon determinations suspected of contamination, those with poor collagen preservation, or those derived from burnt bones were excluded. Reliable radiocarbon dates obtained from bone or charcoal were calibrated using the IntCal20 calibration curve^96^, while those from marine shell samples were calibrated using Marine20^97^, applying a ΔR correction of 0^98^. Chronologies also incorporated luminescence (OSL/TL) and uranium-thorium (U/Th) dates, with associated 1σ uncertainties^99^. The resulting modelled dates, along with their standard deviations, were integrated into the database and used as the temporal framework for analysing species occurrences across sites.

### -Species Distribution Models

In a recent study^100^, we built Species Distribution Models (SDM) for each MIS3 secondary consumer species. This includes: *Ursus arctos, Ursus spelaeus, Panthera spelaea, Panthera pardus, Felis silvestris, Lynx lynx, Lynx pardinus, Crocuta crocuta, Canis lupus, Cuon alpinus, Vulpes vulpes, Vulpes lagopus, Gulo gulo, Meles meles, Martes marteş H. sapiens* and *H. neanderthalensis*. The assignment of archaeological assemblages to either *Homo sapiens* or *Homo neanderthalensis* was based on the following criteria:

a. Assemblage integrity: Assemblages were only included if stratigraphic integrity was clearly supported in the literature. Layers exhibiting evidence of post-depositional mixing or stratigraphic integrity issues were excluded. Justifications for these exclusions are documented on a case-by-case basis in the database.
b. Direct evidence: When human skeletal remains, ancient DNA, or proteomic data unambiguously identified a hominin species associated with a particular archaeological layer, that identification was accepted independently of the absence of associated lithic artifacts.
c. Cultural attribution: In the absence of direct human biological evidence, cultural indicators were used to infer human presence, provided there is strong scholarly consensus on the authorship of specific techno-complexes. Accordingly, assemblages attributed to the Middle Palaeolithic, Mousterian or Micoquian industries were classified as representing *H. neanderthalensis*. In contrast, Upper Palaeolithic cultures such as the Neronian, Aurignacian, Gravettian, Lincombian-Ranisian-Jerzmanowician (LRJ), and Early and Initial Upper Palaeolithic assemblages were used as evidence for the presence of *H. sapiens*.

As part of this study, we applied the same Species Distribution Modelling (SDM) approaches to primary consumer taxa with body masses exceeding 1 kg. The analysis included the following species: *Coelodonta antiquitatis, Mammuthus primigenius, Stephanorius hemitoechus, Ovibos moschatus, Equus hydruntinus, Equus ferus, Bison priscus, Bos primigenius, Saiga tatarica, Megaloceros giganteus, Alces alces, Cervus elaphus, Rangifer tarandus, Capreolus capreolus, Dama dama, Capra ibex, Capra pyrenaica, Capra caucasica, Rupicapra rupicapra, Rupicapra pyrenaica, Sus scrofa, Lepus europaeus, Lepus timidus, Oryctolagus coniculus, Castor fiber*, and *Marmota marmota*.

To build the SDMs, we used the HadleyCM3 model^101^, obtained from the Pastclim R package^102^. HadleyCM3 is a coupled general climate model with active atmosphere, ocean and sea ice components. The spatial resolution is 0.5º * 0.5º, with the 17 climate and 5 environmental variables at 1000-year intervals^101^. We selected this paleoclimate model because its bias-corrected values have been recently validated against paleoclimate data derived from pollen-based temperature and precipitations, as well as stable oxygen values from different stalagmites in Europe^103^.

To minimize site clustering and mitigate potential spatial autocorrelation, we spatially thinned the species occurrence data, ensuring a minimum separation of 0.5º between any two occurrence points. For each species, we delineated a calibration area by constructing a convex hull around all known occurrence points and applying a buffer equivalent to 10% of the maximum distance between those points^104^. This buffered area defined the region from which background (pseudoabsence) points were sampled, ensuring these were drawn from areas environmentally and geographically relevant to the species’ known distribution^105^.

We then extracted climate and environmental variables for all occurrence and background points. Species distribution models (SDMs) were trained using four algorithms: Generalized Linear Models (GLM), Generalized Additive Models (GAM), Maximum Entropy (MAXENT), and Bayesian Additive Regression Trees (BART). For GLM, GAM, and MAXENT, pseudoabsence points were selected by randomly sampling 50 temporally matched locations within the calibration area for each presence record^106^. In the case of BART, we used a balanced dataset with equal numbers of presences and pseudoabsences, repeated the modelling process ten times, and generated an ensemble mean projection from the resulting BART models^107^. For each algorithm, we evaluated the performance with the Area Under the Curve (AUC) and the Boyce Index (BI). Only models achieving an AUC greater than 0.7 were retained for projection, ensuring that poorly calibrated models were excluded^108^.

Since habitat suitability and probability of presence can be influenced by species prevalence, we used habitat favourability values, which correct for prevalence effects^109^. These values express the degree to which a given patch or cell aligns with the species’ ecological requirements and thus offer a standardized basis for interspecific comparison^109,110^. To integrate predictions from different SDM algorithms, we applied a weighted ensemble approach. Specifically, for each species i and grid cell q, the ensemble prediction (WAi) was calculated as a weighted average of the predictions from all SDMs:

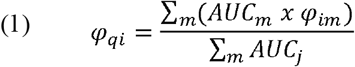

The weighted average favourability prediction (*φ*) is calculated as the sum of predictions for site *q* across *m* individual models (GLM, GAM, MAXENT and BART when the AUC values were larger than 0.7) weighted by their respective AUC values and normalized by the sum of all AUC values^111^.

To select the predictive predictors for each species, we began by removing highly correlated variables (Pearson’s r^2^ ≥ 0.8) from the initial set of 21 climate variables derived from the HadleyCM3^101^ model. Next, we applied a function to identify the optimal set of predictors for each species and SDM algorithm. This function systematically evaluated all possible combinations of the remaining variables using three selection criteria: (1) combinations with any variable exhibiting a Variance Inflation Factor (VIF) above 5 were excluded to prevent multicollinearity; (2) among the valid sets, the one yielding the highest AUC was selected; and (3) when multiple combinations had similar AUC values, the set with the lowest Root Mean Square Error (RMSE) was chosen^100^.

To evaluate model robustness and transferability, we implemented five-fold spatial block cross-validation^111^. Accordingly, the study area was randomly divided into five spatial blocks of roughly equal size, and models were trained and tested five times, each time using one block for testing and the remaining four for training (Supplementary Fig 3). Block size was defined using the cv_spatial_autocor function from the BlockCV R package^112^, which determines appropriate spatial resolution based on autocorrelation patterns. Only models that demonstrated strong predictive accuracy and spatial transferability in this cross-validation process (AUC>0.7) were retained for further analysis.

### -Carrying Capacity of primary consumers

Carrying capacity (CC) is the maximum population size of a given species that an environment can sustain over a specific period^113^. In this study, CC was estimated for each primary and secondary consumer species in each grid cell across Europe at 1,000-year intervals between 55 and 27 kyr BP, based on habitat favourability, total available biomass (TAB), and both intra- and inter-guild competition.

Net Primary Productivity (NPP) was obtained with the Miami model^114^:

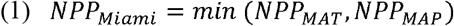

Where:

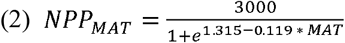

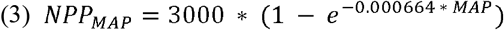

MAT is the mean annual temperature (ºC) and MAP the mean annual precipitation (mm/yr). Both MAT and MAP values were obtained from the HadleyCM3^101^ model. We selected this modelling approach over the BIOM4 model because, although both produce similar NPP trends for the MIS3 period (Supplementary Fig 1), the Miami model yields higher NPP values, making it more suitable for our goal of estimating CC. In addition, MAT and MAP values from the HadleyCM3^101^ model have been validated for this timeframe in Europe based on pollen- and isotope-based reconstructions^18^.

In a previous study^20^, a modelling approach was proposed to estimate herbivore carrying capacity based on the bottom-up effects of food chain regulation driven by NPP, the herbivore guild composition in a specific area or region, and the allometric relationships between body size and population density. Yet, in addition to available biomass, habitat favourability is a critical factor influencing both the presence and abundance of species in a specific area^115^. To incorporate the effect of habitat favourability on herbivore carrying capacity, we adapted the previous modelling approach as detailed below.

The biomass of an herbivore population is obtained by multiplying its population density (D) by the mean adult body mass of the species. Accordingly, the total biomass of all herbivore species (THB) in a given area *q* could be expressed as follows:

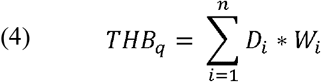

Where *n* is the number of herbivore species in the community, *D* the population density of species *i* expressed as ind/km^2^, and *W* is the mean body mass of both sexes in kg. Damuth^116^ demonstrated that population density changes allometrically with body size:

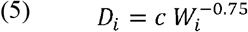

Where *D* is expressed in ind/km^2^, *c* is a constant and *W* is the mean body mass in kg. Thus, *D* can be substituted in equation 5:

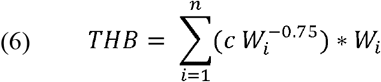

Furthermore, *c* can be estimated as follows:

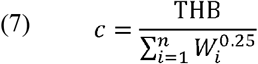

To calculate the c value, it is necessary to first estimate total herbivore biomass (THB). THB that an ecosystem can support is influenced by NPP, so we used a previously proposed predictive equation to estimate THB from NPP^20^:

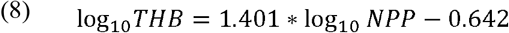

However, instead of using the mean THB, we selected the upper 95th percentile of THB values, thereby focusing on the upper bound of the response distribution (Supplementary Fig. 4). It should be noted that as herbivore populations spread from the core to the outer edges of their geographic ranges, they tend to occupy less favourable habitats and are found at lower densities^115^. To incorporate this effect of differential habitat favourability between primary consumers in each specific cell the distribution of THB among primary consumers was weighted according to the habitat favourability of each species in cell q:

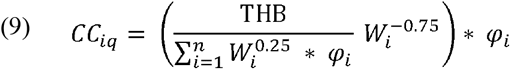

Where CC of species i is expressed as ind/km^2^ and *φ* is the habitat favourability of species i on cell q (Supplementary Fig. 5). Accordingly, THB is allocated across species adjusting for their metabolic requirements, represented through body size exponents, their habitat favourability, and the herbivore guild composition. Body weight for each species was obtained from the Phylacine database^117^, and habitat favourability was obtained from the species distribution models described above (see Species Distribution Models subsection). This modelling approach, previously validated without incorporating habitat favourability and using 674 herbivore density surveys^18^, is here further validated using the Minimum Number of Individuals (MNI) from MIS3 archaeological sites. We found a significant positive correlation between MNI and herbivore carrying capacity (p-value < 0.001, r^2^ 0.2) (Supplementary Table 2).

### -Carrying capacity of secondary consumers

The energy requirement of each secondary consumer species was estimated based on their mean adult body mass, using Farlow’s equation^118^. For human species, however, Resting Metabolic Rate (RMR) was estimated with Kleiber’s equation^119^ and then daily energy requirements were obtained by multiplying RMR by a physical activity level of 1.5^120^. This energy requirement (in kcal/individual) was converted into demanded biomass (dB) in kg per individual per year (kg/yr/ind) using a conversion factor of 1.5 kcal/g^121^. To account for prey size preferences of each secondary consumer species, prey was categorized into four body size classes: small (1-10 kg), medium (10-100 kg), medium-large (100-500 kg), and large (>500 kg). Following previous work ^121–123^, the dB of each species was then adjusted by its specific prey preferences (PP) according to the percentage each species feeds on each prey size category (Supplementary Table 3). Thus, the biomass demanded (dB) of species i is:

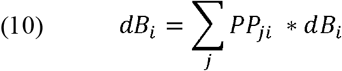

Here, dB is expressed in kg/year and represents the total biomass required by the secondary consumer species i. The variable PP denotes the average percentage of predation exerted by the secondary consumer species i on prey of body size j. PP was obtained from the CarniDIET database^124^ for extant species and from literature for extinct species (Supplementary Table 3). For omnivore species, PP includes only the percentage of their biomass requirement derived from animal matter, so it takes into account the percentage of the diet that relies on both plant and animal resources. That is, dB represents the biomass required by species i based on its energy needs, prey size preferences, and the animal-to-plant composition of its diet. The percentage of the diet composed by animal and plants was obtained from the Phylacine database^117^.

Following the Paleosynecological Model (PSEco) proposed by^123^, the primary consumer species included within the prey preference (PP) spectrum of each secondary consumer species were not estimated based on the mean adult body mass of herbivore species. Instead, it was derived from the distribution of body masses within each primary consumer population^125,126^. The rationale behind this is that a fully-grown, large-sized herbivore may fall outside the preferred prey size range for a medium-sized carnivore; yet, juvenile individuals of that herbivore species, due to their smaller sizes, may still fall within the carnivore’s preferred prey size spectrum^41^. To estimate the population structure of each herbivore species, we computed survival profiles with Weibull models^127–129^ that assume stable and stationary populations. Thus, we used the following stability function (S):

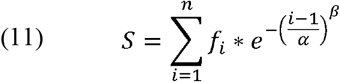

Where n is the total number of age classes of the species, i is the age or time period of the individuals in the cohort, f is the fecundity for each class i, α represents the scale parameter of the Weibull distribution and β the shape parameter of the Weibull distribution. The summation over i from i to n aggregates the survival contributions from all individuals, so in a stable population S should be 1. To ensure S = 1, we need to find appropriate values of α and β. To this aim, we created a loop in R where the code generated a set of random β values for between 0 and 1:

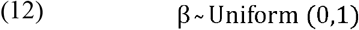

For each β value, it searched an α value using a numerical root-finding method (with the uniroot function) that identifies α values such that S = 1. This procedure was repeated 100 times, and then we used the average value (Supplementary Fig. 6). Once α is estimated, the proportion of individuals alive at each age class is computed as follows:

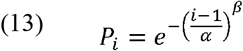

Where P is the probability that an individual survives to age i given the estimated parameters. To compute the total number of individuals by age, we scaled the survival probability to the total population size for the species:

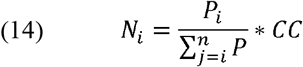

Where N is the number of individuals per km^2^ of age class i, n is the total number of age classes, and CC is the total number of individuals of the species per km^2^. As this provides the number of individuals per age class in a stable population, the number of individuals that do not survive to the next age multiplied by their respective body weights represents the available biomass for secondary consumers that can be obtained without affecting the stability of the population. Thus, the biomass available for each secondary consumer species is:

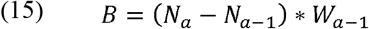

Where *N* represents the number of individuals of age *a* and *W* its body mass. Age-specific body masses were estimated with Gompertz growth curves^130^, which take adult and neonate body masses as inputs ^123,125,126,131^. In addition, to compute the population structure of each herbivore species, information on longevity, age at first reproduction, litter size and litters per year was obtained from the PanTHERIA database^132^ and the Animal Diversity Web for extant species, and from the literature for extinct species (Supplementary Table 4)^133^

We estimated edible biomass (EB) for each grid cell and time step by summing the available herbivore biomass across all size categories and applying a wastage factor to reflect the proportion of prey biomass consumable ^121,123^. The wastage factors used were: 0.8 for small, 0.75 for medium, 0.65 for medium-large, and 0.6 for large-sized herbivores^134^. Therefore, for each cell:

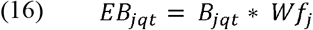

Where EB is the edible biomass of herbivore size class j in cell q at time t, B is the total available biomass, and Wf is the wastage factor for size class j.

As in the herbivore biomass calculation, habitat favourability was used to spatially weight the biomass demand of each secondary consumer species. The spatially-adjusted demanded biomass of species i in cell q is:

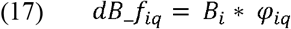

Where dB_f is the annual biomass requirement of species i and *φ* is the habitat favourability of species *i* on cell *q*. A lower habitat favourability reduces both the expected presence and the effective biomass demand of species i in cell q, indirectly increasing the biomass availability for other predators. The available biomass of prey category j for secondary consumer species i in cell q is:

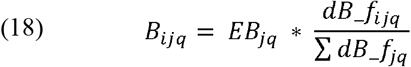

Where EB is the edible biomass for the herbivore weight-size category j on cell q expressed in kg/year. The term 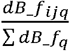 normalizes the proportion demanded by secondary consumer i on prey j, ensuring that the sum B across all species predating on j equals the herbivore biomass of prey category j.

Lastly, the carrying capacity of species i in cell q is estimated by summing the biomass available to them across all prey size classes and dividing by their individual biomass demand:

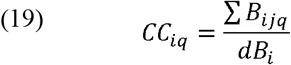

To quantify the intensity of competition among secondary consumers, we used the CC values to compute the Global Competition Index, as proposed by Rodríguez-Gómez et al.^135,136^:

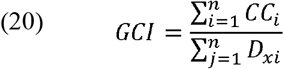

Where CC is the carrying capacity as defined in equation 19, and D_x_ is the mean population density of species i derived from the allometric equations proposed by Damuth^137^ .Thus, this index provides a measure of how closely species approach their theoretical population densities, as predicted in present-day ecosystems. Thus, a GCI < 1 suggests populations are below theoretical expectations in extant ecosystems, indicating low intra-guild competition, while values near 1 reflect populations approaching expected densities based on their body masses in present.

### Agent Based Model

In this study, we built the NEAnderthal Replacement (NEAR) agent-based model to assess the extent to which climate-driven ecological changes contributed to the extinction of *Homo neanderthalensis*. The aim of the NEAR model is not to replicate the historical event, but to explore the combined effects of potential extinction drivers and assess their relative influence. Detailed documentation, following the standardised Overview, Design concepts and Details protocol^138^ is available in Supplementary Note 1.

The environment is a two-dimensional grid composed of 199 * 117 cells, each representing a spatial unit of the European subcontinent (Supplementary Fig 7). There are two types of cells: sea-patches and land-patches. Only land-patches include carrying capacity (CC) values for *H. sapiens* and *H. neanderthalensis*, as described in the “Carrying capacity of secondary consumers” subsection.

Agents represent human bands, each characterised by: 1) species (*H. sapiens* or *H. neanderthalensis*), 2) population size (number of individuals), 3) probability of interbreeding, 4) number of interbreeding events, 5) baseline net growth rate, 6) logistical mobility radius, 7) residential logistical mobility radius, 8) mating territory size, 9) inter-band mobility, 10) carrying capacity, and 11) distance to other agents.

Each time step in the model represents one generation. The simulation runs for 28,000 years, from 55,000 to 27,000 years BP, or until *Homo neanderthalensis* becomes extinct. If *Homo sapiens* populations vanish from Europe during this period, the model registers it as a “failed” colonization attempt. As part of the initial experiment setup, the user specifies the following parameters: 1) whether to simulate one species (*Homo neanderthalensis*) or two species (*H. neanderthalensis* and *H. sapiens*); 2) whether to include Heinrich events, during which the carrying capacity (CC) of each patch randomly decreases by up to 30% during Heinrich stadial 5, 4, and 3^20^; 3) the number of *H. sapiens* arrival waves and the number of bands per wave; 4) the percentage of meat in the agents’ diet; 5) the initial number of agents; 6) the level of demographic stochasticity; 7) the baseline growth rate of agents; 8) the maximum allowable territory overlap between bands; 9) the probabilities of interspecific mating; 10) the logistical and residential mobility of agents; 11) the size of the agents’ mating territory; and 12) the frequency of inter-band mobility among individuals (Supplementary Fig 8).

In each time step, the carrying capacity (K) of each band is determined by considering the carrying capacity (CC) of adjacent habitat patches along with the band’s logistical mobility. This logistical mobility, which is set by the user, represents the average distance that a band travels from its base camp to gather resources. Bands with higher mobility values exploit larger areas. However, if another band is located within this mobility range, the effective K is reduced proportionally to reflect the actual territorial overlap, with the maximum allowed overlap controlled through a user-defined parameter (Supplementary Note 1). Accordingly, while CC denotes the human carrying capacity of each cell, K refers to the carrying capacity specific to each band. Unlike CC, K depends not only on the CC of the cell but also on the band’s mobility and the degree of geographic overlap with other bands. On the other hand, CC and K define a population ceiling, not the actual population size. The actual population size of each Neanderthal and *H. sapiens* band is simulated as follows:

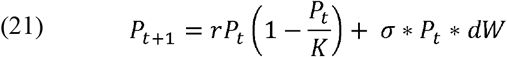

The first term of the equation is the classic Verhulst logistic growth equation^139^. Here, r is the intrinsic baseline growth rate defined by the user, P represents the population size of the band at time t, and K is the specific carrying capacity of the band. Under this deterministic growth dynamic, the population growth rate decreases as P approaches K (Supplementary Figure 9). In contrast, the second term of the equation introduces stochasticity, representing unpredictable fluctuations in growth rates, resource availability, or other ecological factors that may lead to catastrophic mortality events or sudden demographic changes. The parameter σ controls the magnitude of these fluctuations and is specified by the user. Larger values of σ result in greater variability in population size (Supplementary Figure 9). The variable dW represents a Wiener process (or Brownian motion), a noise term drawn from a standard normal distribution. As a result, actual population changes result from a combination of deterministic logistic growth and stochastic variation (Supplementary Figure 9).

Bands approaching their K may undergo fission, during which a subset of individuals emigrate, decreasing the original band’s size (Supplementary Figure 8). If migration occurs, the band searches within its residential mobility radius for patches with equal or greater CC and moves to the best available patch. After relocating, K is recalculated and population dynamics are updated accordingly. The number of emigrants is randomly drawn up to a user-specified maximum percentage. Migrants can either join nearby existing bands with available capacity or establish a new band. In cases where the newly formed band includes individuals from different species, the band’s species identity is assigned according to the majority. Over time, if the composition shifts such that more than 75% of members belong to a different species, the band’s characteristics are updated to reflect the majority species. Each agent also maintains a mating territory, potentially extending beyond its residential mobility, which is specified by the user. Mating can occur both within and between species, contingent on the presence of other bands within this mating radius and the user-defined probability of interspecific mating. The model tracks interbreeding events when such cross-species reproduction happens. Bands that fall below two individuals are deemed extinct and removed from the simulation (Supplementary Note 1).

## Acknowledgements

This research was funded by the European Research Council under the European Union’s Horizon 2020 Research and Innovation Programme (grant agreement number 818299; SUBSILIENCE project; https://www.subsilience.eu). G. R-G holds a postdoctoral position under the “Atracción de Talento Investigador César Nombela” program (Ref. 2023-T1/PHHUM-29222), which is co-funded by the Community of Madrid and the Complutense University of Madrid. We thank all of our colleagues from the EvoAdapta group for constant enriching discussions.

## Author contributions

M.V.-C., A.B.M.-A. and G.R.-G. designed the study. M.V.-C. and G.R.-G. recompiled the data. M.V.-C. run the models and analysed the data. All authors contributed to evaluating the outcomes. M.V.-C. led the writing with critical input from A.B.M.-A. and G.R.-G.

## Data availability

All datasets used in this study are available from OSF (osf.io/4aqjk). Supplementary Information is available for this paper.

## Code availability

All R and Netlogo codes used to perform the analyses reported in this manuscript are available from OSF (osf.io/4aqjk).

