## Supplementary for "Inter-band connectivity and climate shaped Neanderthal extinction and *Homo sapiens*’ dispersal"

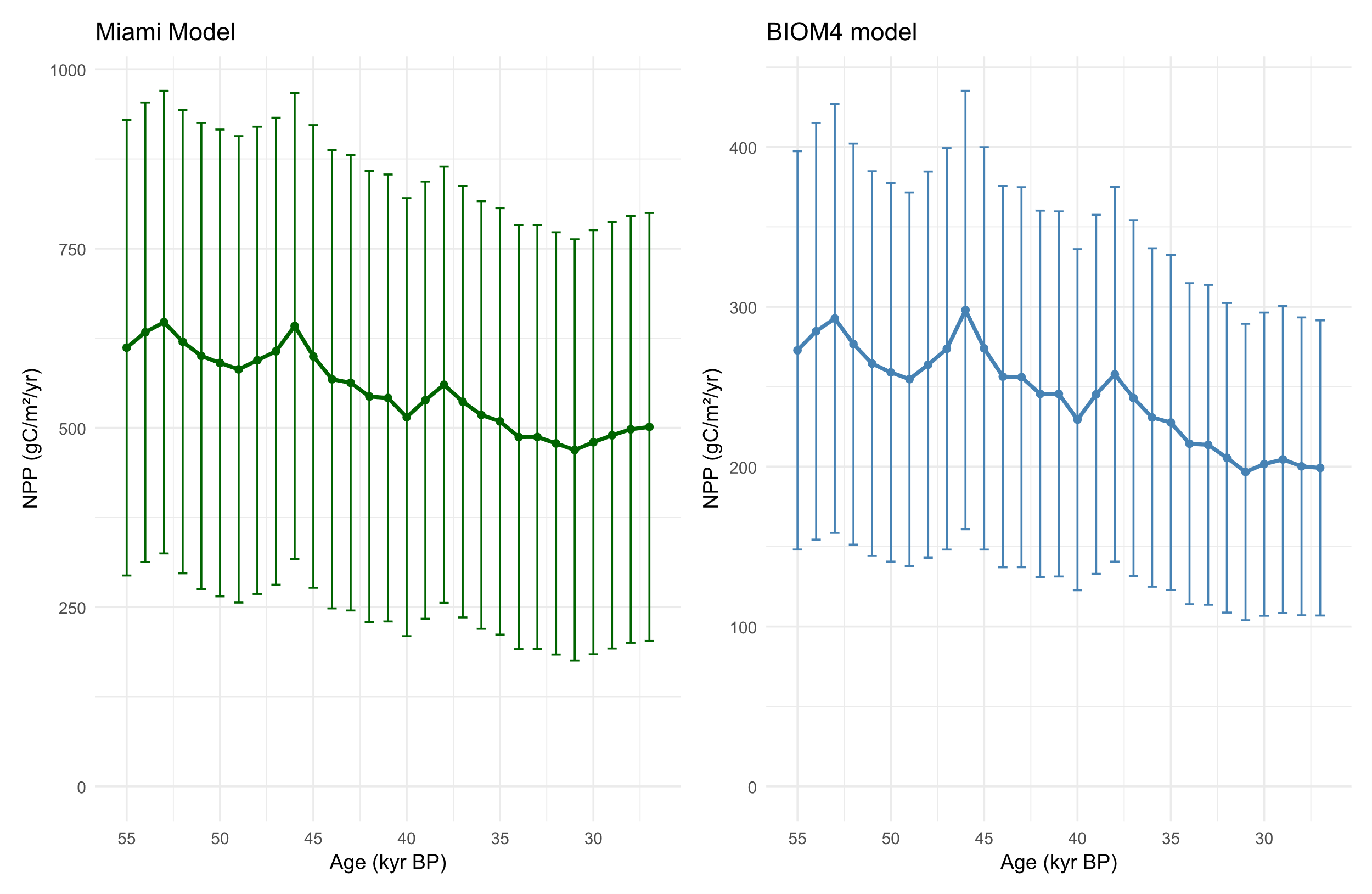

**Supplementary Fig. 1.** Net Primary Productivity during MIS3 according to the Miami model ^2^ (left) and the BIOM4 model ^1^ (right) ; the latter was used in this study. For details, see Methods section.

| Parameter | Description | Default | | Range tested |
| --- | --- | --- | --- | --- |
|  |  | Value | Unit |  |
| n-number | Initial number of Neanderthal bands | 200 | Agents | - |
| s-number | Initial number of *H. sapiens* bands | 200 | Agents | - |
| Two human species | One or two human species | - | - | [true, false] |
| Heinrich events | Heinrich events included | - | - | [true, false] |
| %-meat-in-diet | Percentage of meat included in diet | - | % | [30 10 60] |
| Number of arrival waves | Number of *H. sapiens* arrival waves | 3 | Waves | [1 1 6] |
| Sigma | Demographic stochasticity | 0.4 | % | [0.2 0.1 0.8] |
| Interspecific-mating-prob | Interspecific mating probability | 0.5 | % | [0 0.1 1] |
| Territory-overlap | Percentage of maximum territory overlap between bands | 0.2 | % | [0.1 0.1 0.8] |
| n-r | Neanderthal base net growth rate | 0.1 | % | [0.1 0.1 0.3] |
| s-r | *H. sapiens* base net growth rate | 0.1 | % | [0.1 0.1 0.3] |
| n_logistical_mobility | Neanderthal logistical mobility radius | 15 | Km | [10 5 20] |
| s_logistical_mobility | *H. sapiens* logistical mobility radius | 15 | Km | [10 5 20] |
| n-%inter-band-mobility | Percentage of Neanderthal maximum inter-band mobility | 0.2 | % | [0.1 0.1 0.3] |
| s-%inter-band-mobility | Percentage of *H. sapiens* maximum inter-band mobility | 0.2 | % | [0.1 0.1 0.3] |
| n-territory_size | Neanderthal residential mobility area | 2 | Cells | [1 1 4] |
| s-territory_size | *H. sapiens* residential mobility area | 2 | Cells | [1 1 4] |
| n-mating-territory | Neanderthal mating territory size | 1.5 | Factor | [1 0.5 2] |
| n-mating-territory | *H. sapiens* mating territory size | 1.5 | Factor | [1 0.5 2] |
| wave-pop | Number of individuals per each *H. sapiens* arrival wave | 100 | Individuals | [50 50 200] |
| **Supplementary Table 1.** Parameters included in the NEAR agent-based model and values tested. | | | | |

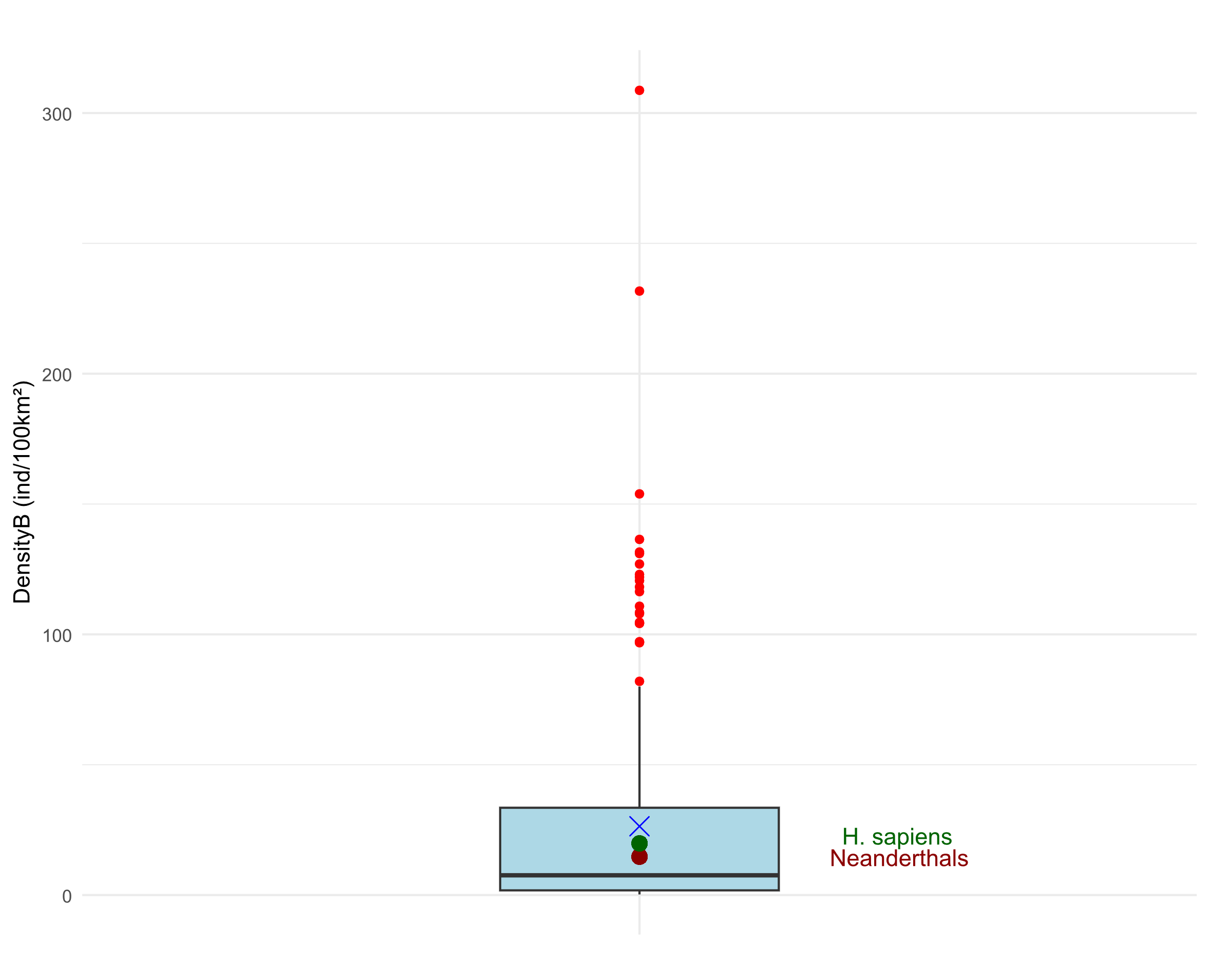

**Supplementary Fig. 2.** Box and whisker plot with the population density of 215 contemporary hunter-gatherer societies based on data provided by Binford^3,4^. The centre line of the box-plot corresponds to the median, the blue X to the mean value, the box limits to the first and third quartiles, the whiskers to 1.5x interquartile range and the red dots to the outliers. The green and garnet points indicate the estimated carrying capacity for *H. sapiens* and *H. neanderthalensis* respectively.

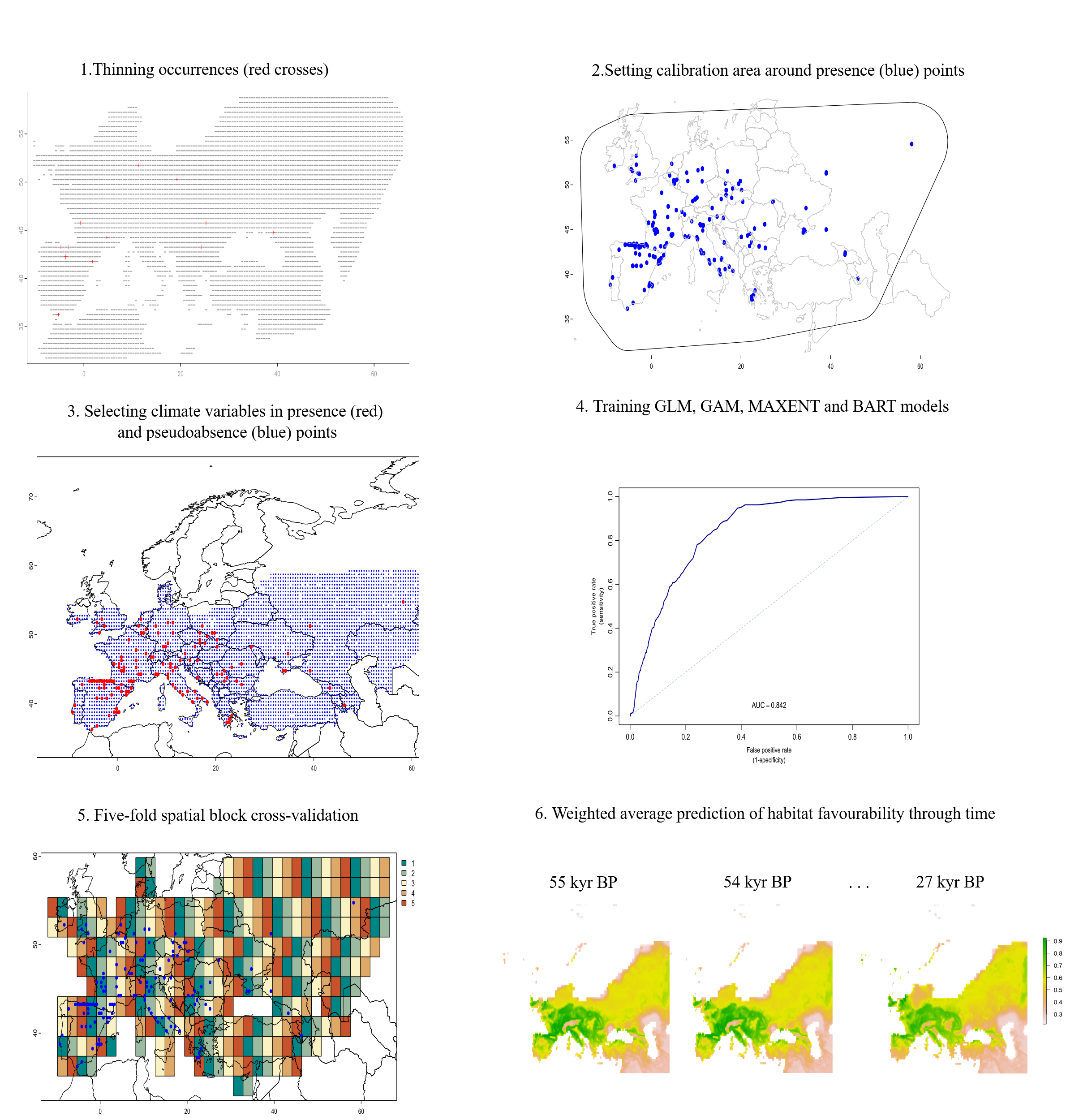

**Supplementary Fig. 3.** Workflow summarising the main steps used to build SDMs for all primary and secondary consumer species. For details, see Methods section.

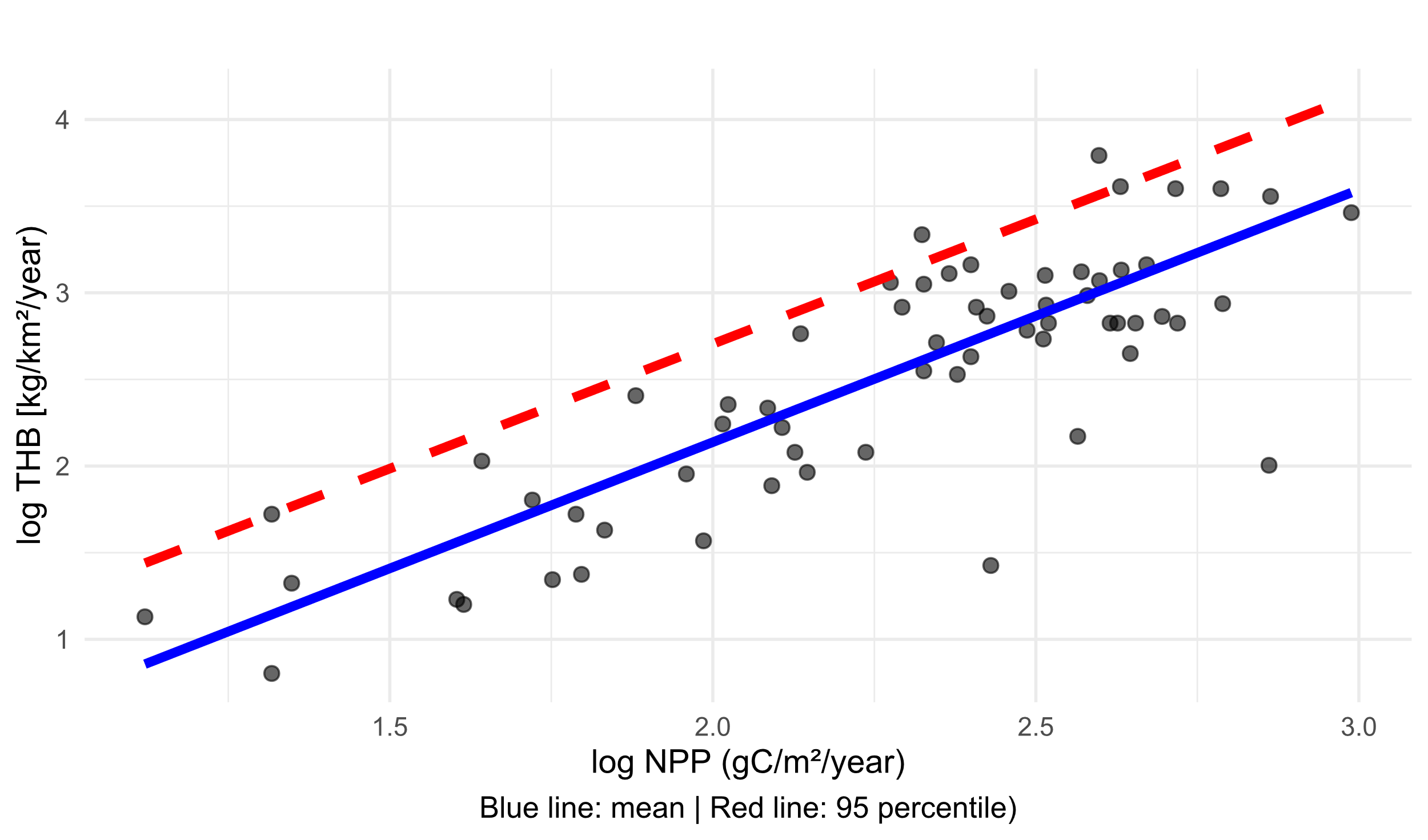

**Supplementary Fig. 4**. Relationship between log-transformed Net Primary Productivity (log NPP, gC/m²/year) and log-transformed Total Herbivore Biomass (log THB, kg/km²/year) in present-day ecosystems. The blue solid line indicates the mean trend used in previous studies to compute herbivore biomass^5,6^, while the red dashed line shows the 95th percentile, highlighting the upper bound of the observed distribution, which was used in the current study to compute herbivore carrying capacity.

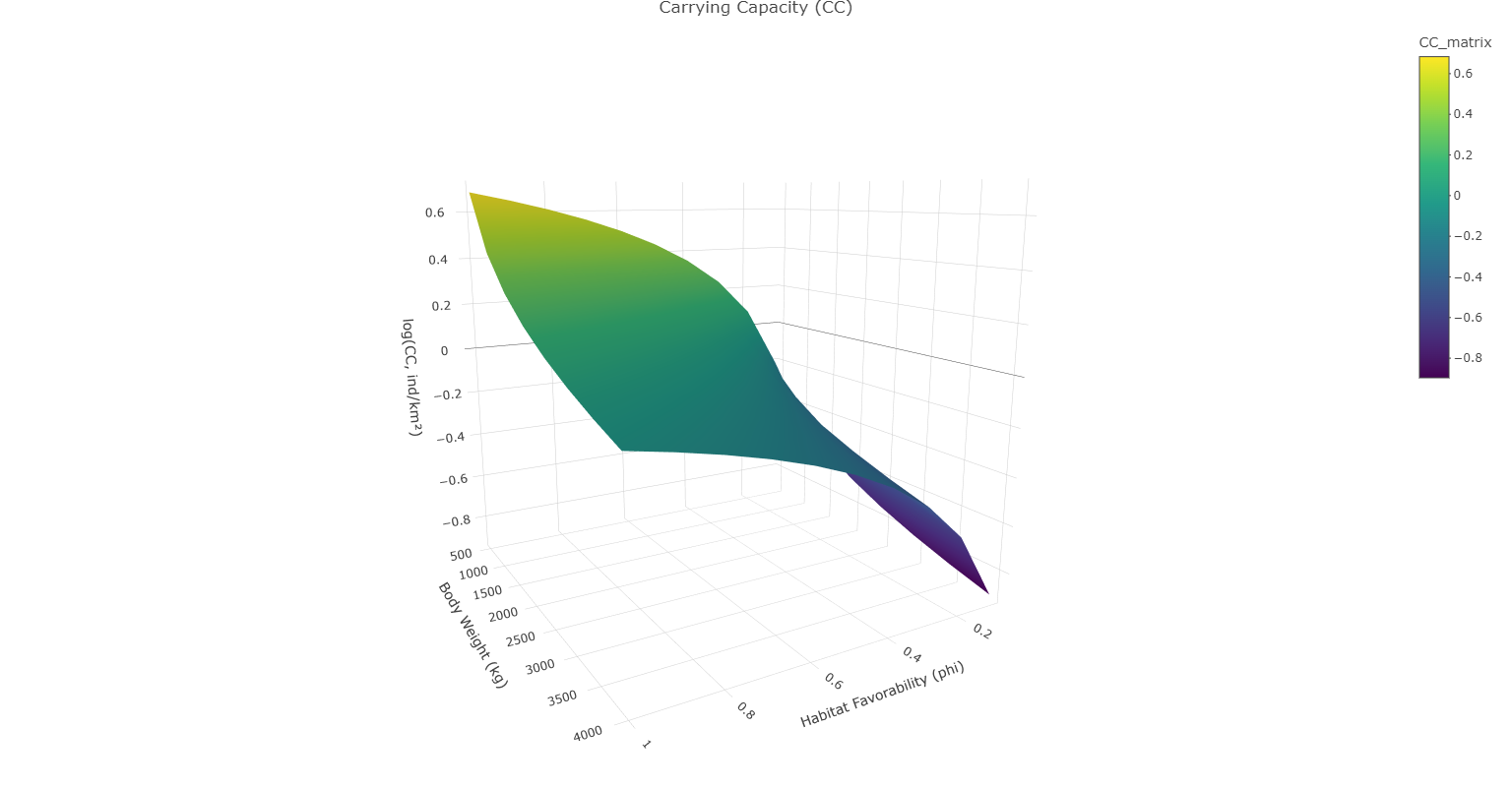

**Supplementary Fig. 5**. Response of carrying capacity (CC) to varying values of body mass and habitat favourability as independent variables, based on the methodological approach proposed in this study (see Equation 9 in the Methods section). The plot shows the influence of body size on CC, considering all possible habitat favourability values for each body weight.

|  | Value | Std. Error | DF | t-value | p-value | r2 | AIC | BIC |
| --- | --- | --- | --- | --- | --- | --- | --- | --- |
| (Intercept) | 0.3836 | 0.0263 | 480 | 14.60 | <0.001 | 0.2 | 699.36 | 716.81 |
| logDENS | 0.1074 | 0.0301 | 480 | 3.57 | 0.0004 |  |  |  |
| **Supplementary Table 2.** Statistic values showing a positive correlation between the estimated herbivore carrying capacity (logDENS) and the minimum number of individuals (MNI) recovered from the archaeological record. | | | | | | | | |

| Species | Adult Body Mass (kg) | Energy requirements (kcal/day) | % Meat in diet | Prey preference spectrum (%) | | | | Ref |
| --- | --- | --- | --- | --- | --- | --- | --- | --- |
|  |  |  |  | Small (1-10 kg) | Medium (10-100 kg) | Medium-large (100-500 kg) | Large (>500 kg) |  |
| *Ursus spelaeus* | 390.416 | 14927.32 | 1.00 | 25.0 | 25.0 | 25.0 | 25.0 | ^7–10^ |
| *Ursus arctos* | 180.52 | 8720.32 | 10.00 | 25.0 | 25.0 | 25.0 | 25.0 | ^7,8,10^ |
| *Panthera spelaea* | 380.189 | 14653.74 | 100.00 | 0.0 | 43.0 | 45.0 | 12.0 | ^7,8^ |
| *Panthera pardus* | 54.99 | 3808.76 | 100.00 | 2.0 | 80.0 | 18.0 | 0.0 | ^7,8^ |
| *Crocuta crocuta* | 62.99 | 4186.88 | 98.50 | 0.0 | 45.2 | 40.1 | 14.7 | ^7,8^ |
| *Canis lupus* | 32.18 | 2621.96 | 100.00 | 3.0 | 30.0 | 64.0 | 3.0 | ^7,8^ |
| *Meles meles* | 13 | 1394.16 | 5.00 | 100.0 | 0.0 | 0.0 | 0.0 | ^7,8,11^ |
| *Vulpes vulpes* | 5.31 | 747.03 | 10.00 | 100.0 | 0.0 | 0.0 | 0.0 | ^7,8,11^ |
| *Vulpes lagopus* | 4.86 | 702.33 | 10.00 | 50.0 | 50.0 | 0.0 | 0.0 | ^7,8,11^ |
| *Felis silvestris* | 5.5 | 765.56 | 5.00 | 100.0 | 0.0 | 0.0 | 0.0 | ^7,8,11^ |
| *Gulo gulo* | 17.01 | 1681.43 | 80.00 | 75.0 | 18.8 | 6.3 | 0.0 | ^7,12^ |
| *Lynx lynx* | 17.95 | 1745.65 | 100.00 | 65.0 | 35.0 | 0.0 | 0.0 | ^7^ |
| *Lynx pardinus* | 9.4 | 1112.20 | 100.00 | 75.0 | 25.0 | 0.0 | 0.0 | ^7^ |
| *Cuon alpinus* | 12.76 | 1376.17 | 90.00 | 22.2 | 44.4 | 27.8 | 5.6 | ^7^ |
| *Martes martes* | 1.3 | 280.19 | 60.00 | 83.3 | 16.7 | 0.0 | 0.0 | ^7^ |
| *Homo neanderthalensis* | 76 | 2702.71 | [30-60] | 14.0 | 46.0 | 30.0 | 10.0 | ^7^ |
| *Homo sapiens* | 59.5 | 2249.45 | [30-60] | 14.0 | 46.0 | 30.0 | 10.0 | ^7^ |
| **Supplementary Table 3.** Body size, energy requirements, and prey spectra according to prey weight-size categories for each secondary consumer species. | | | | | | | | |

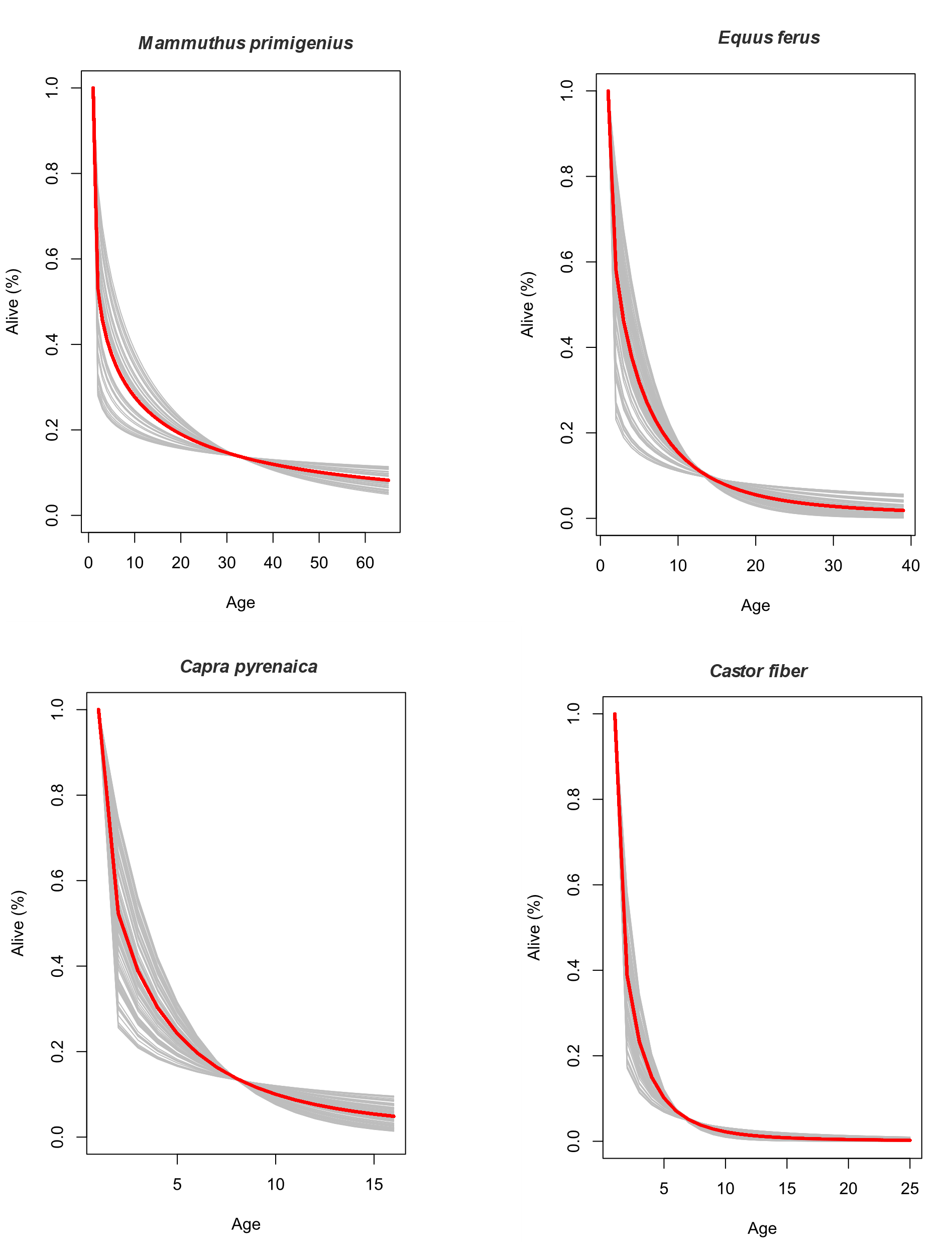

**Supplementary Fig. 6.** Simulated population structures (n = 100; grey lines) for four species based on fitted Weibull models. Red lines denote the mean structure derived from the simulations.

| Species | Adult Body Mass (kg) | Neonate Body Mass (kg) | Age at First Birth (d) | Litter Size | Litters Per Year | Longevity (m) | Ref |
| --- | --- | --- | --- | --- | --- | --- | --- |
| *Mammuthus primigenius* | 6000.00 | 101.00 | 11.25 | 1.13 | 0.24 | 65.00 | ^7,8,13^ |
| *Stephanorius hemiothecus* | 3287.05 | 6.75 | 1.00 | 0.36 | 41.75 | 47.00 | ^7,8,13^ |
| *Coelodonta antiquitatis* | 2692.55 | 41.75 | 6.75 | 1.00 | 0.36 | 47.00 | ^7,8,13^ |
| *Bison priscus* | 950.00 | 39.48 | 2.62 | 1.00 | 0.91 | 25.00 | ^7,8,13^ |
| *Bos primigenius* | 900.00 | 37.67 | 2.62 | 1.00 | 0.91 | 25.00 | ^7,8,13^ |
| *Megaloceros giganteus* | 699.84 | 29.78 | 2.86 | 1.00 | 1.10 | 20.80 | ^7,8,13^ |
| *Alces alces* | 357.00 | 13.00 | 3.33 | 1.20 | 1.00 | 27.00 | ^7,8,13,14^ |
| *Ovibos moschatus* | 340.50 | 10.67 | 3.50 | 1.01 | 0.89 | 24.00 | ^7,13^ |
| *Equus hydruntinus* | 227.13 | 28.73 | 3.50 | 1.00 | 0.67 | 38.80 | ^7,8,13^ |
| *Equus ferus* | 200.00 | 28.63 | 3.50 | 1.00 | 0.67 | 38.80 | ^7,8,13^ |
| *Cervus elaphus* | 131.25 | 82.56 | 2.72 | 1.09 | 0.90 | 26.80 | ^7,8,13^ |
| *Sus scrofa* | 101.05 | 0.81 | 0.85 | 4.52 | 1.50 | 21.00 | ^7,13,14^ |
| *Rangifer tarandus* | 86.03 | 5.49 | 2.69 | 2.00 | 1.10 | 20.20 | ^7,13,14^ |
| *Capra ibex* | 85.17 | 2.78 | 2.50 | 1.11 | 0.89 | 22.30 | ^7,13,14^ |
| *Dama dama* | 56.25 | 4.70 | 3.00 | 1.00 | 1.10 | 25.00 | ^7,13,14^ |
| *Capra pyrenaica* | 50.00 | 2.90 | 3.00 | 1.10 | 1.10 | 16.00 | ^7,13,15,16^ |
| *Saiga tatarica* | 29.00 | 3.50 | 1.32 | 1.50 | 1.10 | 12.00 | ^7,13^ |
| *Rupicapra rupicapra* | 26.10 | 2.25 | 3.75 | 1.00 | 1.10 | 22.00 | ^7,13^ |
| *Capreolus capreolus* | 22.50 | 1.21 | 2.00 | 1.79 | 1.10 | 17.00 | ^7,13^ |
| *Castor fiber* | 19.00 | 0.55 | 2.50 | 2.95 | 1.00 | 25.00 | ^7,13^ |
| *Lepus europaeus* | 3.74 | 0.12 | 0.69 | 2.14 | 4.40 | 12.00 | ^7,13^ |
| *Lepus timidus* | 3.05 | 0.11 | 0.87 | 3.16 | 3.00 | 18.00 | ^7,13^ |
| *Marmota marmota* | 2.01 | 0.03 | 2.36 | 4.00 | 1.00 | 18.00 | ^7,13^ |
| *Oryctolagus coniculus* | 1.96 | 0.04 | 0.33 | 5.24 | 4.50 | 18.00 | ^7,13^ |
| **Supplementary Table 4.** Life history traits of each primary consumer species included in this study and used to compute age-specific body masses with Gompertz growth curves and population structure with Weibull models. | | | | | | | |

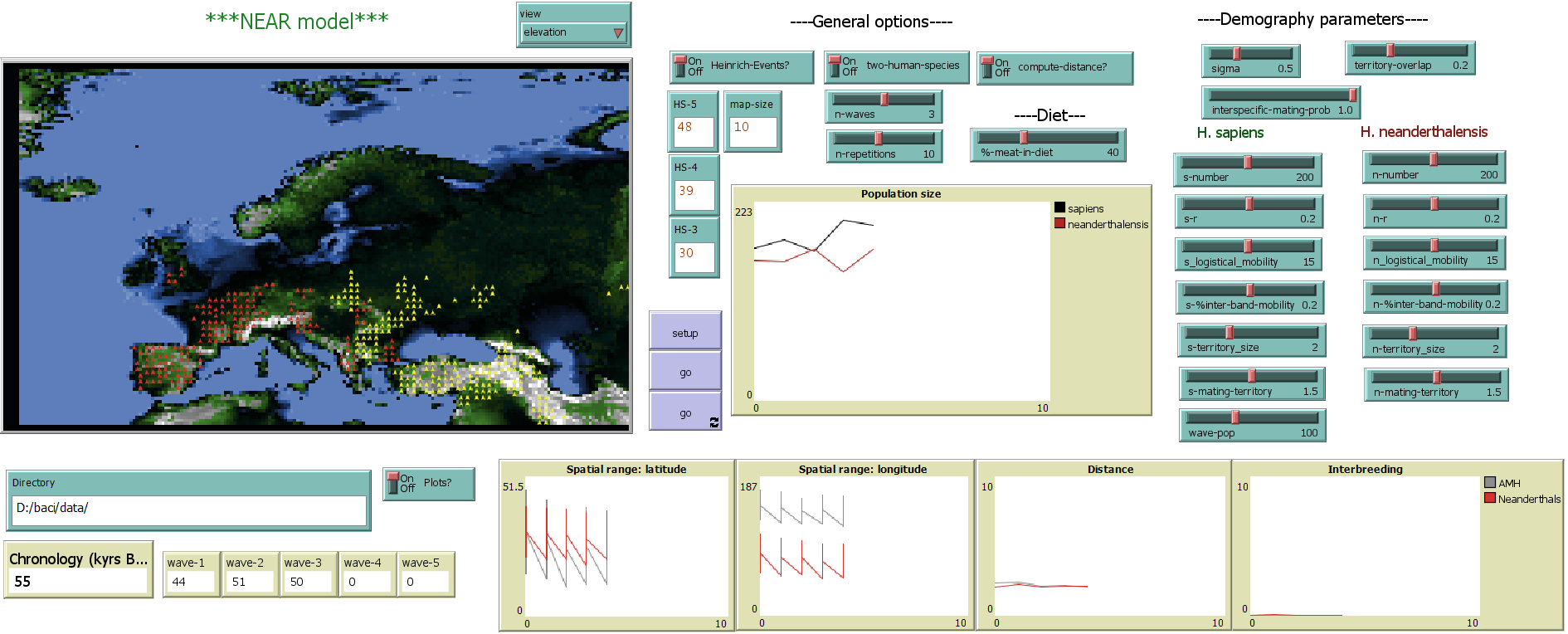

**Supplementary Fig. 7**. NEAR model interface in NetLogo with controls and sliders, spatial view, and graphs tracking model data.

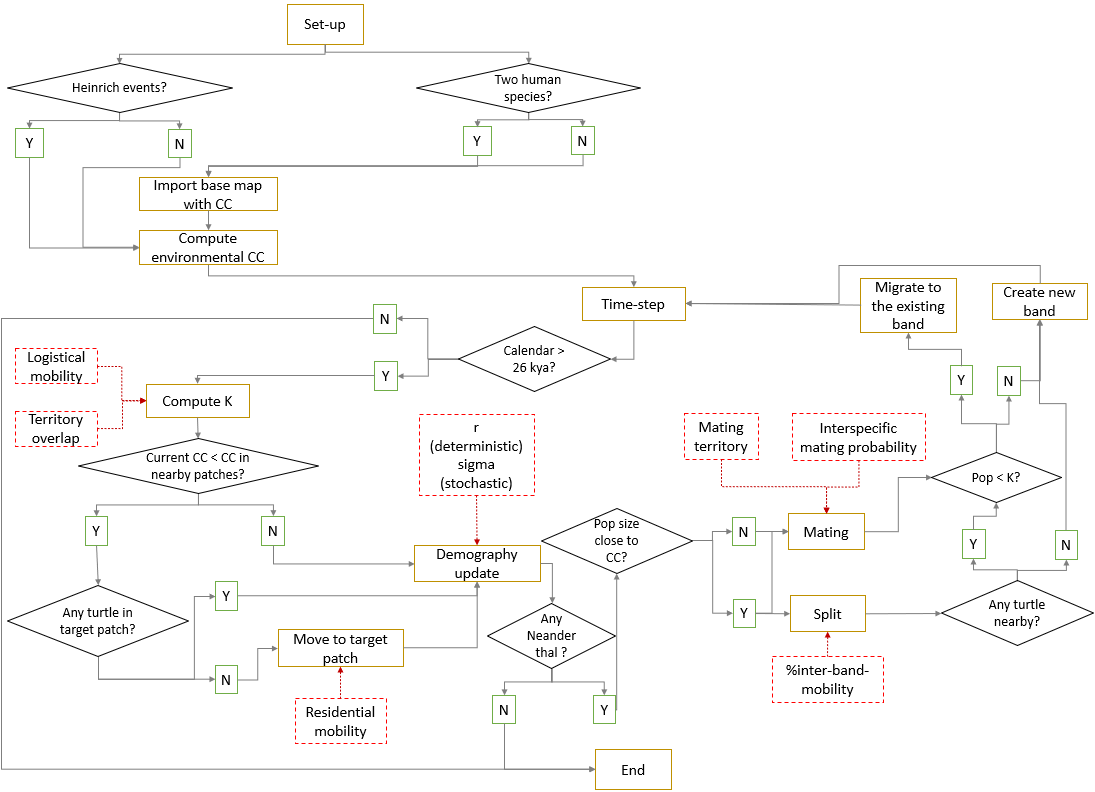

**Supplementary Fig. 8**. Model schedule overview. Flow chart of the model showing the sequence of procedures (rounded cells) and checks (diamonds) the model follows each time step. Dashed lines indicate the main variables set by the user.

| 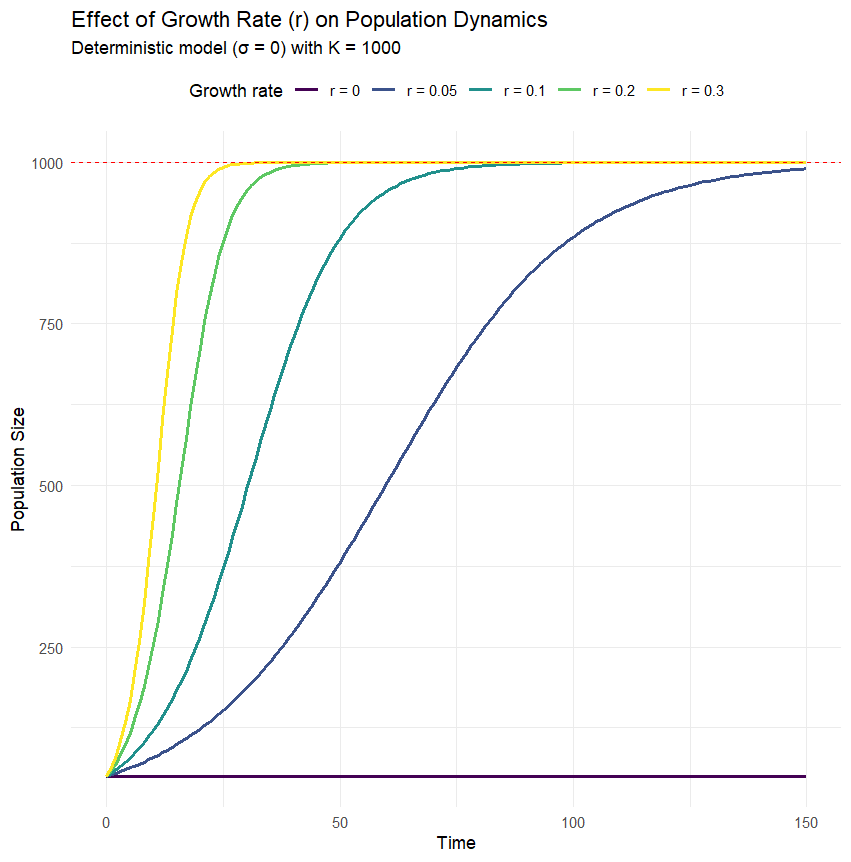 | 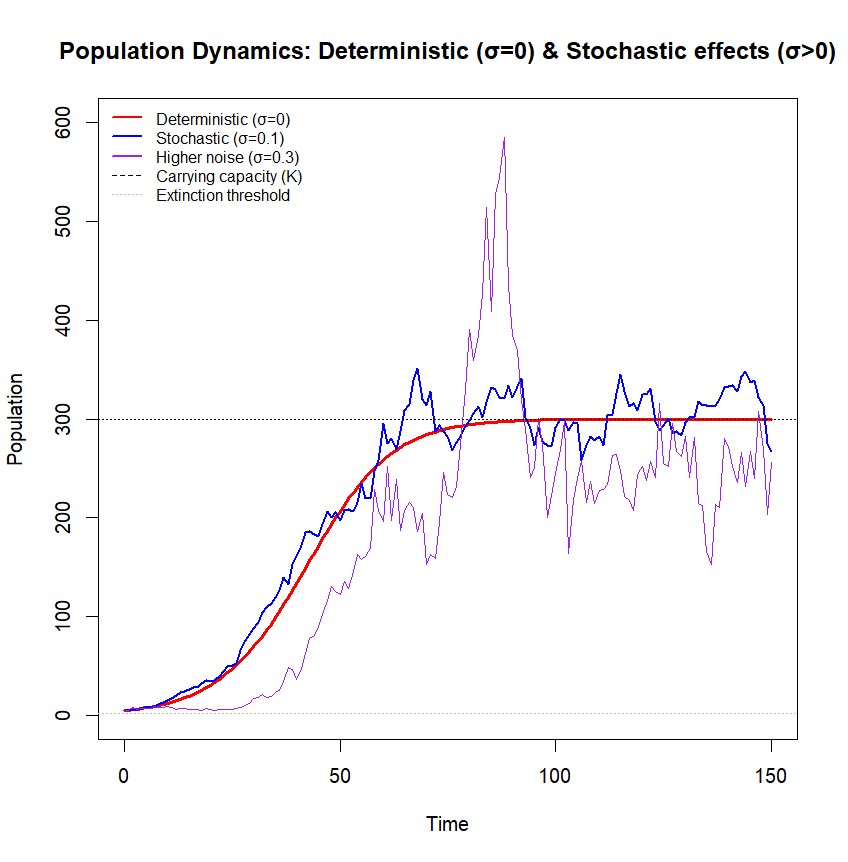 |
| --- | --- |
| **Supplementary Fig. 9.** Deterministic (right) and stochastic (left) dynamics of population growth under varying values of baseline growth rate (r) and environmental stochasticity (σ), as defined in Equation 21 of the Methods section. Horizontal dashed lines indicate the carrying capacity. | |

**Supplementary Note 1**

**Overview, Design concepts, and Details (ODD) protocol.**

**NEAR: the Neanderthal Replacement model**

**Purpose**

Agent-based models are computational tools that simulate the interactions of autonomous agents within a defined environment, enabling researchers to explore how complex dynamics and emergent patterns arise from specific ecological or behavioural rules^17–19^. The Neanderthal Replacement model (NEAR) is an agent-based model designed to provide a tool to identify and analyse the main factors that contributed to the replacement of Neanderthals by *Homo sapiens* in Europe. The model is specifically designed to test two hypotheses:

1. Climate-driven changes in carrying capacity triggered the extinction of *H. neanderthalensis* in Europe.

2. The arrival and expansion of *H. sapiens* in Europe had a greater impact on Neanderthal extinction than climate changes.

To this aim, the model evaluates how extinction risk responds to four key factors: (a) proportion of meat intake, (b) baseline population growth rate, (c) demographic stochasticity, and (d) mobility patterns, including logistical, residential, and inter-band mobility. These dynamics are simulated under two distinct scenarios: (1) one where *Homo sapiens* is absent, (2) and another where *H. sapiens* arrives in Europe (Supplementary Table 1). In the two-species scenario, the model further incorporated additional variables to assess their impact on Neanderthal extinction risk and timing, including the number of *H. sapiens* migration waves, their population size, the probability of interspecific mating, and the extent of territorial overlap or segregation between the two human species.

**Agent classes, variables and temporal and spatial scales**

The environment is a two-dimensional grid composed of 199 * 117 cells, each representing a spatial unit of the European subcontinent, with the size of each cell defined by the user (Supplementary Fig. 7). There are two types of cells: sea-patches and land-patches. Only land-patches include carrying capacity (CC) values for *H. sapiens* and *H. neanderthalensis*. These CC values are computed for each cell at 1,000-year intervals during MIS3, based on: human, prey and predators’ habitat favourability; Net Primary Productivity; available biomass of prey; energetic requirements, prey size preferences, and intra-guild competition (for details, see Carrying Capacity subsection in Methods section).

Agents represent human bands, each characterised by: 1) species (*H. sapiens* or *H. neanderthalensis*), 2) band size (number of individuals), 3) interbreeding probability, 4) number of interbreeding events, 5) baseline growth rate, 6) logistical mobility, 7) residential logistical mobility or territory size, 8) mating territory size, 9) percentage of inter-band mobility, 10) carrying capacity, and 11) distance to other agents (Supplementary Table 1).

**Process overview and scheduling**

Each time step in the model represents one generation. The user defines how many generations occur per 1,000 years. The simulation runs for 28,000 years, from 55,000 to 27,000 years BP, or until *Homo neanderthalensis* becomes extinct. If *Homo sapiens* populations vanish from Europe during this period, the model registers it as a failed colonization attempt. If Heinrich events are included (via the "Heinrich-Events" toggle), the carrying capacity (CC) of land patches will randomly decrease by up to 30% during Heinrich Stadial 5, 4, and 3^6^.

In each time step, a band’s carrying capacity (K) is calculated based on the CC of surrounding patches and the band’s logistical mobility. Logistical mobility, defined by the user, represents the average distance (in km) a band travels from its base camp to acquire resources. Higher mobility values reflect exploitation of larger areas. However, if another human band is within this mobility radius, K is proportionally reduced based on territorial overlap. The maximum allowable overlap is also user-defined (via the “territory-overlap” setting).

As a band’s population approaches its CC, its probability of migrating increases, following a rule based on the population-to-CC ratio (Supplementary Fig. 8). If migration occurs, the band searches for a better patch within its mobility radius, defined as residential mobility, and will relocate if the new patch offers an equal or higher CC (Supplementary Fig. 8). After migration, CC is recalculated and demographic dynamics are updated.

Population growth is influenced by both deterministic logistic growth and stochastic fluctuations (for details, see Methods section). Bands near their CC may split, and mating can occur either within or between species, potentially forming new bands. When a band splits, a portion of its members emigrate, reducing the original band's size. The number of migrants is randomly determined, with a user-defined maximum (%inter-band-mobility). Migrants may either join existing nearby bands (if those have available CC) or form a new band if more than five individuals remain unabsorbed. If the new band consists of mixed species, its identity is determined by the species with the majority. Over time, if more than 75% of the members are from a different species than the original, the band adopts the attributes of the majority species.

Each agent also has a mating territory, which may extend beyond its logistical mobility and is defined by the user. Mating can occur between same or different species, depending on the presence of bands within this radius and the interspecific mating probability (also user-defined). When interspecies mating and reproduction occur, the model records an interbreeding event. A band with fewer than two individuals is considered extinct and removed from the simulation (Supplementary Fig. 8).

**Design concepts**

The NEAR model incorporates spatially explicit estimations of human carrying capacity (CC) to explore extinction risks under varying specific factors (i.e., independent variables), including:

a) The presence of one (*H. neanderthalensis*) or two (*H. neanderthalensis* and *H. sapiens*) human species in Europe;

b) The percentage of meat included in human diet;

c) Mobility strategies, including logistical mobility, residential mobility, and mating territory;

d) Probability of interbreeding;

e) The extent of territorial overlap or segregation between bands.

The following basic principles have been applied in the NEAR model:

- Agents represent human bands that evolve demographically (through internal population growth, splitting, and mixing), migrate, and may go extinct.
- Agents do not anticipate environmental change; they react to fluctuations in human carrying capacity (CC) by adjusting their population sizes and/or migrating.
- Each run serves as an experiment, where all variables are held constant except the one being tested (i.e., one of the independent variables listed above, following the One-Factor-At-a-Time approach described in the Statistical analyses subsection).

Adaptation: Human bands adapt to their environment by exploiting available meat resources and adjusting their population sizes. If the human CC cannot support the current population, bands may split and/or migrate.

Objectives: Agents aim solely to increase their population size or migrate to areas where their band sizes can be sustained. To achieve this, they require a CC sufficient to support their populations, as well as social or spatial structures (bands within a certain radius) that facilitate reproduction.

Prediction: Agents do not explicitly predict future environmental conditions. They do not possess memory or learning mechanisms related to environmental cycles. However, they do: a) record the number of interspecific breeding events, b) track the proportion of individuals of each species within the band, c) estimate the CC of potential new locations before migrating.

Interaction: Agents interact with: a) the environment by exploiting resources, b) other bands both directly (e.g., through exchange or absorption of individuals) and indirectly (e.g., overlapping foraging areas reduce CC proportionately).

Sensing: Agents know the human CC within their residential mobility area, detect the presence of other agents in the area, and evaluate the population pressure of neighbouring agents when deciding to split or migrate.

Stochasticity. Human CC is spatially distributed based on available biomass after taking into account habitat favourability of prey and predators, fluctuations in Net Primary Productivity and intra-guild competition dynamics among secondary consumers for each 1,000 years during MIS3. However, CC defines a population ceiling, not the actual band size. Actual demographic evolution incorporates a stochastic component (sigma) that represents random fluctuations. The magnitude and frequency of this stochasticity are user-defined via the “sigma” button (Supplementary Fig. 9).

Observations: the following data is collected from the model: Timing and occurrence of the extinction of *H. neanderthalensis* or *H. sapiens* in Europe, mean and standard deviation of the longitude and latitude occupied by agents every 1,000 years, number of agents, number of interbreeding events.

**Initialization**

The model world is represented as a grid matrix derived from a set of ASCII files, each containing the CC values for *Homo sapiens* and *H. neanderthalensis* at 1,000-year intervals during MIS3 in Europe. Before running the model setup, these files must be placed in different folders according to the percentage of meat included in the agents’ diet, and the path to this directory must be specified in the “Directory” window (Supplementary Figure 7).

The cell size of the grid is determined by the map resolution, which is set by the user via the “map-size” box. Additionally, an elevation raster file for Europe is loaded. Following the methodology of Gravel-Miguel and Wren^20^, the elevation of each cell is interpolated using the bicubic_2 method.

During setup: 1) Agents (human bands) are randomly distributed on land patches where CC > 0.001 individuals/km²; 2) each band starts with a population size between 20 and 30 individuals; 3) *H. neanderthalensis* bands are distributed across all suitable land-patches in Europe, while *H. sapiens* bands are initially placed only in eastern Europe; 4) to aid visualization, the residential mobility area of each agent is displayed as a circle around the agent during setup.

**Input**

The NEAR model uses a series of raster files (.asc) as input data, representing human carrying capacity for each cell/patch at 1,000-year intervals between 55,000- and 27,000-years BP. Human CC for each species is estimated based on the following factors:

a) Fluctuations in the base of the food chain, specifically Net Primary Productivity (NPP)

b) Changes in habitat favourability for each prey species in each cell during MIS3

c) Variations in available prey biomass within each cell during MIS3

d) Changes in habitat favourability for each predator species in each cell during MIS3

e) Energetic requirements and prey size preferences of secondary consumers, including humans, within each cell during MIS3.

For further details on the carrying capacity models, see the Carrying Capacity subsection in Methods.

**Submodels**

Habitat favourability of each prey and predator, including humans, was obtained with four Species Distribution Models’ algorithms: Generalized Linear Models (GLM), Generalized Additive Models (GAM), Maximum Entropy (MAXENT), and Bayesian Additive Regression Trees (BART). For each algorithm, we evaluated the performance with the Area Under the Curve (AUC) and the Boyce Index (BI). Only models achieving an AUC greater than 0.7 were retained for projection, ensuring that poorly calibrated models were excluded ^21^. The paleoclimate data to build these SDMs were obtained from the HadleyCM3 model^22^, since its bias-corrected values have been recently validated against paleoclimate data derived from pollen-based temperature and precipitations, as well as stable oxygen values from different stalagmites in Europe^23^. The outputs of these SDMs consists in one layer with the habitat favourability of each species in Europe at 1000-year intervals between 55 and 27 kya. For details, see Species Distribution Model Subsection in Methods.

Net Primary Productivity (NPP) was estimated with the Miami model. ^2^ Mean annual temperature (°C) and precipitation (mm/year) for each cell/patch were obtained from the HadleyCM3 ^22^ model and used as input data. Based on NPP, the herbivore guild composition, their body masses and habitat favourability, we developed a model to estimate the carrying capacity of each herbivore species in each cell. For details, see Carrying Capacity of primary consumers subsection in Methods.

Carrying capacity of secondary consumers was estimated using the Paleosynecological Model (PSEco)^24^. The maximum abundance of each species, including humans, was computed based on energetic requirements, prey size preferences, habitat favourability, and the CC of primary consumers, categorised into four weight classes (1-10 kg; 10-100 kg; 100-500 kg; >500 kg). Prey preference spectrum of each secondary consumer was not estimated based on the mean adult body mass of herbivore species, but from the distribution of body masses within each primary consumer population, as juveniles often fall within a carnivore’s preferred prey range even if adults do not. Thus, we estimated herbivore population structures using Weibull survival models, and then estimated the CC of each secondary consumer species by dividing the biomass demanded by all carnivore species in each cell by the available herbivore biomass in the same cell (for details, see Carrying Capacity of secondary consumers subsection in Methods)

The carrying capacity of each cell sets an upper limit on the population size but does not directly determine the actual population of each band. The number of individuals in each band is modelled by the following equation:

$$P_{t+1}=rP_{t}\left( 1-\frac{P_{t}}{K} \right)+ \sigma*P_{t}*dW$$

In the first term, P_t_ represents the current population size of the band, 𝑟 is the intrinsic net growth rate defined by the user, and K is the human carrying capacity. The value of K depends on the carrying capacity of the cell, the residential mobility of the band, and the degree of geographic overlap with other bands. This term models logistic population growth, where the growth rate slows as the population approaches the carrying capacity K (Supplementary Fig. 9). The second term introduces stochasticity to the model, accounting for random fluctuations in birth and death rates, meat resource availability, and other ecological factors. Here, sigma controls the magnitude of these random fluctuations and is specified by the user. Larger values of sigma lead to greater variability in population size (Supplementary Fig. 9). Accordingly, the number of individuals in each band is computed according to a deterministic and a stochastic component.

**Calibration of the model, testing outputs and statistical analyses**

The NEAR model may generate a large volume of dynamic and high-dimensional data, which makes them difficult to analyse. In this study, we have use One Factor at Time (OAF) to carry out the analyses. Thus, multiple runs were performed for different parameter values at discrete intervals. For details of all values explored, see (Supplementary Table 1). Accordingly, in this study we explored a total of 1,408 parameter combinations (as listed in Table 1). For each combination, the model was run 10 times to account for stochastic variation, which resulted in a total of 14,080 simulations.

To investigate the factors influencing Neanderthal extinction risk, we employed a Classification and Regression Tree (CART) model. The dependent variable was a binary outcome indicating extinction status: extinct (0) or not extinct (1). The model included the following independent variables:

1) Presence or absence of *Homo sapiens*
2) Population size of *Homo sapiens*
3) Number of failed attempts by *H. sapiens* to colonize Europe
4) Number of interbreeding events
5) Inclusion or exclusion of Heinrich events
6) Proportion of meat in the diet
7) Demographic stochasticity
8) Probability of interbreeding
9) Degree of territorial overlap between bands
10) Base net growth rate of Neanderthals
11) Base net growth rate of *H. sapiens*
12) Difference in net growth rate between the two human species
13) Logistical mobility of Neanderthals
14) Logistical mobility of *H. sapiens*
15) Residential mobility of Neanderthals
16) Residential mobility of *H. sapiens*
17) Inter-band mobility
18) Mean distance between Neanderthal bands
19) Mean distance between *H. sapiens* bands
20) Number of arrival waves of *H. sapiens*
21) Mating territory size of Neanderthals
22) Mating territory size of *H. sapiens*

To analyse the correlation between all dependent and independent variables, exploratory plots were constructed, along with the calculation of Spearman correlation coefficients. All data files containing simulation outputs, as well as the R scripts used for the statistical analyses, are available in osf.io/4aqjk
